# Elevated CO2, Warming and Drought Differentially Impact Reproductive and Vegetative Economic Traits in Two Grassland Species

**DOI:** 10.1101/2025.05.15.654190

**Authors:** Murugash Manavalan, Dinesh Thakur, Andreas Schaumberger, Michael Bahn, Zuzana Münzbergová

## Abstract

**Background and Aims:** Since the Industrial Revolution, rising atmospheric CO□, warming, and more frequent droughts have significantly impacted ecosystems. While the response of leaf functional traits to these climate change factors have been widely studied, reproductive traits remain relatively understudied, despite their key role in the diversification and distribution of flowering plants. Here, we investigated how elevated CO□, warming, drought, and their interactions affect floral, leaf and seed traits in two model grassland species. We also examined how these factors influence trait coordination.

**Methods:** Two common grassland species, *Lotus corniculatus* and *Crepis capillaris*, were sampled from a 10-year climate manipulation experiment. We measured resource economic traits related to organ size, construction cost, and dry matter content in both leaves and flowers, along with seed size and number. Univariate and multivariate analyses were used to assess trait responses, and rank-abundance curves were employed to visualize changes in trait coordination across treatments.

**Key Results:** Trait responses to climate change factors varied between species. Drought emerged as the most influential factor, affecting only leaf traits in *L. corniculatus*, but impacting leaf, floral, and seed traits in *C. capillaris*. Across both species, climate change conditions increased leaf construction costs and reduced flower size. In addition, it led to larger leaves in *L.* corniculatus, and fewer seeds in *C. capillaris*. Under extreme climate change conditions, trait coordination became stronger in both species, although *C. capillaris* showed no coordination response specifically to drought.

**Conclusion:** Our results show that floral economic traits, like leaf traits, are responsive to individual and combined effects of climate change factors. This highlights their importance in shaping plant strategies under environmental stress and emphasizes the need to better integrate floral traits into the whole-plant economic framework.

## Introduction

Human activities since the Industrial Revolution have significantly increased greenhouse gas emissions, leading to today’s high levels of atmospheric CO_2_, which is the main driver of the ongoing climate change (Borghi *et al*., 2019; IPCC, 2022). The direct and indirect effects of this change have resulted in rapid alternations of functioning of plants and entire natural ecosystems (Feehan, Harley and van Minnen, 2009). Plants form the foundation of natural ecosystems, serving as primary producers that sustain all other life forms by driving energy flow and nutrient cycling. Due to the significant impact of climate change on ecosystems, it becomes crucial to understand the effects of different climate change factors such as CO_2_, warming and drought on plants (Descamps, Quinet and Jacquemart, 2021). Such knowledge can help to predict and mitigate the adverse effects of climate change on biodiversity and ecosystem services, ensuring the sustainability and resilience of natural ecosystems.

Plant functional traits have long been used by ecologists as useful proxies of plant ecological strategies (Wright *et al*., 2004; Reich, 2014) to gain insights into species response to changing environmental conditions (Liu *et al*., 2024; Tyagi and Kumar, 2024). Recent major climate change factors, particularly elevated CO_2_, temperature, and drought, significantly impact plant traits and physiology across multiple scales, although to different extents (Lukac *et al*., 2010; Albert *et al*., 2011; Bjorkman *et al*., 2018; Abdelhakim, Zhou and Ottosen, 2022). Drought, due to its ease of implementation in experimental studies, seems to be the most extensively studied of all the climate change factors. Among the climatic factors, it produces the most uniform and predictable negative response of range of plant functional traits, including vegetative traits related to competitive ability (plant height and leaf area) (Lidon, 2012; Burkle and Runyon, 2016; Zhou *et al*., 2018; Rodríguez-Alarcón, Tamme and Carmona, 2022), resource use strategy traits such as specific leaf area and specific root length (Jung *et al*., 2014; Wellstein *et al*., 2017; Rodríguez-Alarcón, Tamme and Carmona, 2022) and reproductive traits such as flower size and number, floral rewards, and seed set (Kuppler *et al*., 2021; Akter and Klečka, 2022; Höfer *et al*., 2023).

In contrast, the response of plant traits to factors such as temperature and CO_2_ are more inconsistent and context dependent, varying by species, traits and environmental conditions. While warming has been shown to have significant negative effects on both vegetative and reproductive traits (Niu *et al*., 2000; Liang, 2016; Descamps, Quinet and Jacquemart, 2021), studies have revealed that traits such as leaf area, stem length, stem diameter, fine-root biomass and pollen germination are optimized at certain temperature (Nicotra *et al*., 2008; Dostálek, Rokaya and Münzbergová, 2020; Wang *et al*., 2021; Shin, Yeon and Kim, 2023; Kumarathunge *et al*., 2024) and decline with both temperature increase and decrease. Due to the variability of temperature-optima across species, the effects of temperature on plants are much more species-specific compared to the more consistent effects of drought.

Elevated CO_2_ generally has a positive effect on plants, enhancing photosynthesis, leaf area, carbohydrate concentration, and boosting plant biomass when water is sufficient (Duan *et al*., 2018; Temme *et al*., 2019). Long-term Free-Air CO_2_ Enrichment (FACE) experiments have further validated these findings, showing sustained increases in photosynthetic rates and biomass accumulation over several growing seasons (Ainsworth and Long, 2005; Leakey *et al*., 2009). However, the effects of elevated CO_2_ on plant reproductive traits are far less studied compared to its effects on vegetative traits (but see Wang, Taub and Jablonski, 2015; Glenny, Runyon and Burkle, 2018).

While numerous studies have examined the individual effects of temperature, moisture, and elevated CO- on plant traits, these environmental factors often interact in complex and non-additive ways. This has led to a growing emphasis on investigating their combined effects, particularly on leaf morphology (e.g., Qaderi, Kurepin and Reid, 2006; Albert *et al*., 2011; Mueller *et al*., 2022). However, the interactive impacts of these climate change drivers on reproductive traits—especially in relation to resource allocation strategies—remain largely understudied.

Understanding how plants allocate resources under stress is central to predicting their survival and reproductive success in changing environments. The concept of plant economic spectra provides a valuable framework for this, linking traits such as leaf thickness, tissue density, and dry matter content to construction costs and resource-use strategies (Wright *et al*., 2004; Freschet *et al*., 2010; Díaz *et al*., 2016). While the leaf economic spectrum has been widely studied (Wright *et al*., 2004; Reich, 2014), recent work has extended this framework to reproductive structures through the emerging concept of the floral economic spectrum (Roddy *et al*., 2021; E-Vojtkó *et al*., 2022). This approach identifies key trade-offs in floral trait construction and function, offering insights into how plants balance investment in reproduction with environmental constraints.

Studying floral economic traits under climate change conditions is particularly important, as flowers are critical for reproductive success and pollinator interactions (Ceulemans *et al*., 2017). Environmental stresses such as drought and warming have been shown to affect flower size as well as nectar production and composition, which in turn influence pollinator visitation patterns (Descamps, Quinet and Jacquemart, 2021; Martén-Rodríguez *et al*., 2025). These stresses may also affect the energy and carbon resources that plants allocate in building and maintaining floral tissues, often reflected in shifts related to construction cost-related traits. By examining how floral economic traits respond to environmental stressors, we can better understand the adaptive strategies plants use to optimize reproductive investment and anticipate how plant–pollinator dynamics and ecosystem functions may shift under future climate scenarios.

Most studies examining the response of plant traits to environmental factors focus on the variation of traits in individual plant organs (e.g. Nie *et al*., 2013; Jung *et al*., 2014; Bongers *et al*., 2017; Hai *et al*., 2023; Fenollosa *et al*., 2024). However, different organs respond to environmental factors in distinct ways, and their responses may also covary. For example, an analysis of leaf and root traits across 24 species found that leaf traits exhibited stronger and more homogenous response to drought, while root trait responses were highly variable and species-specific (Lozano *et al*., 2020). This highlights the need to investigate how traits across different organs coordinate under stress. While floral traits are often considered decoupled from leaf traits due to differing selective pressures (Salguero-Gómez *et al*., 2016; E-Vojtkó *et al*., 2022), emerging evidence suggests that coordination does occur. For instance, recent findings show correlations between floral traits such as pollen grain number and nectar volume with leaf traits such as leaf area and leaf dry matter content, indicating that trait integration across organs may be more common than previously assumed and warrants further investigation (Fantinato *et al*., 2025).

The present study seeks to fill the above identified gaps in our knowledge by investigating how climate change factors, namely CO_2_, temperature and drought, affect floral, leaf and seed traits and the covariation between them in two model grassland species. Specifically, this study addresses the following questions and hypotheses.

Q1. How do floral, leaf and seed traits vary in response to elevated CO_2_, warming and drought?

H1. Drought will induce the most consistent and negative changes across plant traits, whereas elevated CO is expected to have more positive effects due to enhanced photosynthetic activity.

Q2. How do these climate change factors interact to affect floral, leaf and seed traits?

H2. In the interaction between temperature and CO_2_, CO_2_ would negate the negative effects of warming. Increasing temperature would exacerbates the negative effects of drought which would be partially mitigated by elevated CO_2_.

Q3. How do climate change factors influence the covariation of flower, leaf and seed traits?

H3. Elevated climate change factors exert a strong filtering action on the covariation of plant traits across different organs, leading to coordinated trait adjustments to adapt to stressful conditions. Similar to leaf traits, floral traits would exhibit adaptive changes in response to individual and combined climate change factors, and that these changes would be strongly coordinated with leaf and seed trait responses.

## Materials and Methods

### Study Site and Experimental Setup

This study was conducted within the framework of a long-term multifactorial climate change experiment (named ClimGrass) at the Agricultural Research and Education Centre (AREC) in Raumberg – Gumpenstein (Meeran *et al*., 2021; Reinthaler *et al*., 2021; Radolinski *et al*., 2025). This study site is located in Styria, Austria (47°29′37″N, 14°06′0″E), 710m above sea level and was established in 2012 in a managed sub-montane C3 grassland. The soil is classified as Cambisol with loamy texture. Aboveground biomass is mown and removed three times a year and plots are regularly bolstered with mineral fertilizer with a total load of 90 kg N ha^-1^y^-1^, 65 kg P ha^-1^y^-1^ and 170 kg K ha^-1^y^-1^. This is done to compensate for the macronutrients lost due to harvests (Pötsch *et al*., 2019). The experimental design comprises of a total of 54 plots (4 × 4m each) representing individual and combined effects of three levels of temperature (ambient, +1.5°C, +3°C), atmospheric CO_2_ concentration (ambient, +150ppm, +300ppm) and drought (yes/no) (Piepho *et al*., 2017; Pötsch *et al*., 2019). The CO_2_ and temperature treatments started in May 2014 while the drought treatments started in May 2017. The different temperature and CO_2_ levels were established based on potential future climate scenarios and were simulated through a combination of infrared heating systems to increase air temperature and a mini-FACE (Free Air CO_2_ Enrichment) approach for fumigation with CO_2_ (Piepho *et al*., 2017; Pötsch *et al*., 2019). Dynamic rainout shelters controlled by rain sensors were deployed on 12 plots to generate water stress in combination with ambient and future climate change. These shelters were only used to restrict rain and induce drought during the summer seasons (Exact dates provided in Supplementary Information Table S8.). Half of the drought plots were maintained at ambient conditions with the remaining half being exposed to the extremes of CO_2_ (+300ppm) and temperature(+3°C) (Radolinski *et al*., 2025; Tissink *et al*., 2025). The plots which acted as control were not heated or fumigated and were equipped with non-functional heaters and/or mini-FACE rings of same shape and size which blew ambient air to account for potential disturbances (Piepho *et al*., 2017; Pötsch *et al*., 2019)

### Plant Sampling and Trait Measurements

For this study, we selected a set of plots representing six treatments: Control (C0T0D0), elevated CO_2_ concentration (+300ppm)(C1T0D0), increased temperature/warming (+3°C)(C0T1D0), drought (C0T0D1), combination of elevated CO_2_ and warming (C1T1D0) and finally combination of elevated CO_2_, warming and drought (C1T1D1). Each of these treatments were represented by atleast 4 plots, providing consistent replication across the manipulated climate conditions. The plots were additionally narrowed down based on the presence of either of two model grassland species, *Lotus corniculatus* L. and *Crepis capillaris* (L.) Wallr. These two species are the two most common insect pollinated species occurring in the experimental plots and have the same flowering period.

*Lotus corniculatus* L. (Fabaceae) is a polycarpic perennial non-clonal herb widely distributed across Europe (Chytrý *et al*., 2021). It flowers in a umbel inflorescence type (Chytrý *et al*., 2021). The corolla is zygomorphic, yellow and consists of five petals each: the upstanding ‘standard’, the lateral two wings and the final two lower petals uniting to form the ‘keel’ with the overall shape of the corolla being butterfly-like (*Nature Gate*, 2011). The plant is primarily insect pollinated.

*Crepis capillaris* (L.) Wallr. (Asteraceae) is an annual herb and is native in Europe being widespread as far north as Denmark and southern Sweden (Stace, 2010; Govaerts, 2024). It flowers in a synflorescence corymbiform inflorescence type. Each flower head contains 50-70 flowers. The corolla is ligulate, 6.0-15.0 mm long, and yellow with a ligule tinged red (Kilian, 2024). It is pollinated by insects but is also capable of selfing (Chytrý *et al*., 2021).

The sampling was conducted on 22^nd^ and 23^rd^ of July 2023. For *L. corniculatus*, a total of 39 individuals were sampled from 28 plots across the 6 treatments, with 5 to 11 individuals sampled per treatment. From each individual, 1–4 flowers and 1–2 leaves were collected. For *C. capillaris*, 45 individuals were sampled from a total of 21 plots with 6 to 10 individuals per treatment. From each *C. capillaris* individual, 1–2 flowers and leaves were sampled (exact number of individuals and plots sampled for each species and treatment have been detailed in Supplementary Information Table S1). The variation in the number of individuals, leaves and flowers is given by their availability in the different treatments (selecting only fully developed, but not wilted and undamaged leaves and flowers/inflorescences). Within each plot, samples were collected at least 10 cm apart with maximum 2 samples per plot and wrapped in water-saturated paper towels and stored in the refrigerator until further processing (approximately 48 hours). Alongside flowers and leaves of both species, seeds of *C. capillaris* were also sampled, dried in paper bags and stored. Developed seeds were not available in *L. corniculatus* and thus could not be sampled.

To estimate Leaf Area (LA), Specific Leaf Area (SLA) and Leaf Dry Matter Content (LDMC), each leaf which was previously wrapped in wet paper towels for at least 48 hours (during sampling) was weighed to determine its saturated mass and then subsequently scanned in a flatbed scanner to determine the Leaf Area (LA). The leaves were dried at 70°C for 72 hours and after which their dry mass was also taken. Leaf area (LA) was estimated using ImageJ software. Specific Leaf Area (SLA) was calculated as Leaf Area (LA) divided by leaf dry mass and Leaf Dry Matter Content (LDMC) was estimated as the ratio of leaf dry mass to saturated mass (Pérez-Harguindeguy *et al*., 2013).

The flowers of the two species were processed differently due to variation in their morphology. The petals of *L. corniculatus* were dissected out from the flowers and weighed to obtain their saturated mass. Afterwards, they were separated out into their individual petal types: one standard petal, two wings and one keel. The separated petals were then flattened and scanned, dried similarly to leaves and then weighed again using a Mettler Toledo Microbalance (d = 0.5μg) to get their dry mass. The scanned petals were processed through ImageJ to obtain their individual petal area. As the keel is composed of 2 fused petals, its area was multiplied by 2. The total petal area of all the petals was taken as the Display Area (DA). The total Specific Petal Area (SPA) of the whole corolla of each flower was calculated (Display Area / Dry Mass). Petal Dry Matter Content (PDMC) was calculated by dividing the total dry mass of the petals by their saturated mass (similarly to leaves the petals have been wrapped in wet tissue for at least 48 hours prior estimating saturated mass).

For *C. capillaris*, 5 individual ray florets were dissected out and then weighed, flattened and scanned to get their fresh mass and petal area. They were then dried and weighed to get their dry mass. Due to the homogeneity of ray florets, the florets were processed together, and their Specific Petal Area (SPA) was calculated as the total area of 5 ray florets divided by their total dry mass. Here, Display Area (DA) was calculated as an average of the 5 individual ray florets. Petal Dry Matter Content (PDMC) was also obtained. The number of seeds in each flowerhead was counted. Seeds number per flower head (SN) was calculated by dividing the total number of seeds taken from a plant by the number of flower heads. The total weight of the seeds in a flower head were weighed using the microbalance mentioned above. The Seed Mass (SM) was then calculated by dividing the weight of all the seeds in a flower head by the number of seeds.

### Statistical Analyses

All statistical analyses were carried out in R software version 4.3.1. AI assistance was used to generate initial R code for certain analytical approaches. Two separate datasets were prepared for the univariate and multivariate analyses. For the univariate analysis, trait values from leaves or flowers of the same individual were retained as separate data points. In contrast, for the multivariate analysis, trait values were averaged per individual prior to analysis. Prior to multivariate analyses, all traits were standardized to a mean of zero and a standard deviation of one to account for differences in measurement scales across traits.

To assess the broad patterns of response of different functional traits to climate change factors, a Redundancy Analysis (RDA) was first performed using the vegan package in R(version 2.6.10; Oksanen *et al*., 2025) followed by a Permutational Analysis of Variance (PERMANOVA) using 9999 permutations to test the significance of the climate change factors in the RDA.

To assess the impact of the climate change factors on individual plant traits, we conducted a set of univariate analyses where first we performed Shapiro-Wilk test to evaluate normality of the trait values. Based on the p-values obtained, the normality of the original trait value was compared with its logarithmic and square-root transformations. The form of the trait value with the highest p-value was selected for inclusion in the linear models, as it indicated the closest approximation to a normal distribution. In *L. corniculatus*, Specific Petal Area and Leaf Dry Matter Content were square-root and log-transformed respectively while all other variables were used in their original form. In *C. capillaris*, only Display Area was kept in its original form with Specific Petal Area, Petal Dry Matter Content, Leaf Dry Matter Content and Seed Number being square-root transformed, and Leaf Area, Specific Leaf Area and Seed Mass being log-transformed. Linear models were used to analyse the effects of CO- (C), temperature (T), drought (D), and their interactions—CO- × temperature (C×T) and (CO- + temperature) × drought (CT×D)—on leaf and reproductive traits of both species separately. All environmental variables and their interactions were included simultaneously as fixed predictors within a single model, without applying variable selection procedures such as stepwise or backward elimination. The “ggplot2” package (v 3.5.1)(Wickham, 2016) was used for creating the plots.

To understand general patterns of covariation across all plant traits, Principal Component Analysis (PCA) was conducted and visualized using the “factoextra” package (v 1.0.7)(Kassambara and Mundt, 2020) in R.

Pair-wise correlations between individual plant traits were analysed by means of a trait correlation network constructed using significant Pearson correlations. A threshold of *r* > 0.2 marked pairwise correlations that were significant at *p* < 0.05. All correlations below this threshold were set to zero, yielding the adjacency matrix A = [*a*_i,j_] with *a*_i,j_ [ [0,1]. Additionally, network connections between any pair of traits are weighted by the absolute correlation strength, |*r*_ij_| (Langfelder and Horvath, 2008; Kleyer *et al*., 2019). Tests such as the Principal Component Analysis (PCA) and network analysis were primarily conducted to under the covariation and network connections between generally unknown traits such as Specific Petal Area (SPA) and Petal Dry Matter Content (PDMC) with well-established traits such as Specific Leaf Area (SLA) and Leaf Dry Matter Content (LDMC).

To assess the covariation of plant traits in response to elevated CO-, increased temperature, and drought, separate Principal Component Analyses (PCAs) were conducted for each treatment. Because the number of individuals per treatment was unbalanced, we performed multiple independent tests using repeated random sampling to standardize the dataset. In each iteration, we randomly selected five individuals per treatment for *L. corniculatus* and six individuals per treatment for *C. capillaris*, ensuring equal sample sizes across treatments. This repeated subsampling approach allowed us to assess the robustness of trait covariation patterns across treatments. The significance of the resulting PCAs and the contribution of individual traits to the significant axes were evaluated using the “PCAtest” package (v 0.0.2)(Camargo, 2025). The proportion of variance explained by the PCA axes for each species was visualized as rank-abundance curves using the ggplot2 package (Wickham, 2016).

## Results

The results are presented in three sub-sections. The first section details the responses of traits of both plant species to the individual and interactive effects of climate change factors. For each species, we first report findings from the multivariate analysis, followed by results from the univariate analyses. The second section examines the covariation among plant traits, while the final section presents how trait covariation is influenced by climate change factors.

### Response of Plant Functional Traits to Climate change Factors

#### Lotus corniculatus

The three climate change factors and their interactions collectively explained 29% of the trait variation in the multivariate redundancy analysis (RDA) (p = 0.002 for the overall model). When tested individually, drought (D) and warming (T) had significant effects, CT×D had marginally significant effects (Fig. 1) while elevated CO_2_ and C×T did not have any significant effect. Among the significant predictors, drought accounted for the highest variance (10.2 %), followed by warming (6.3%) and then CT×D (4.8%) (Supplementary Information Table S2).

**Fig. 1.**
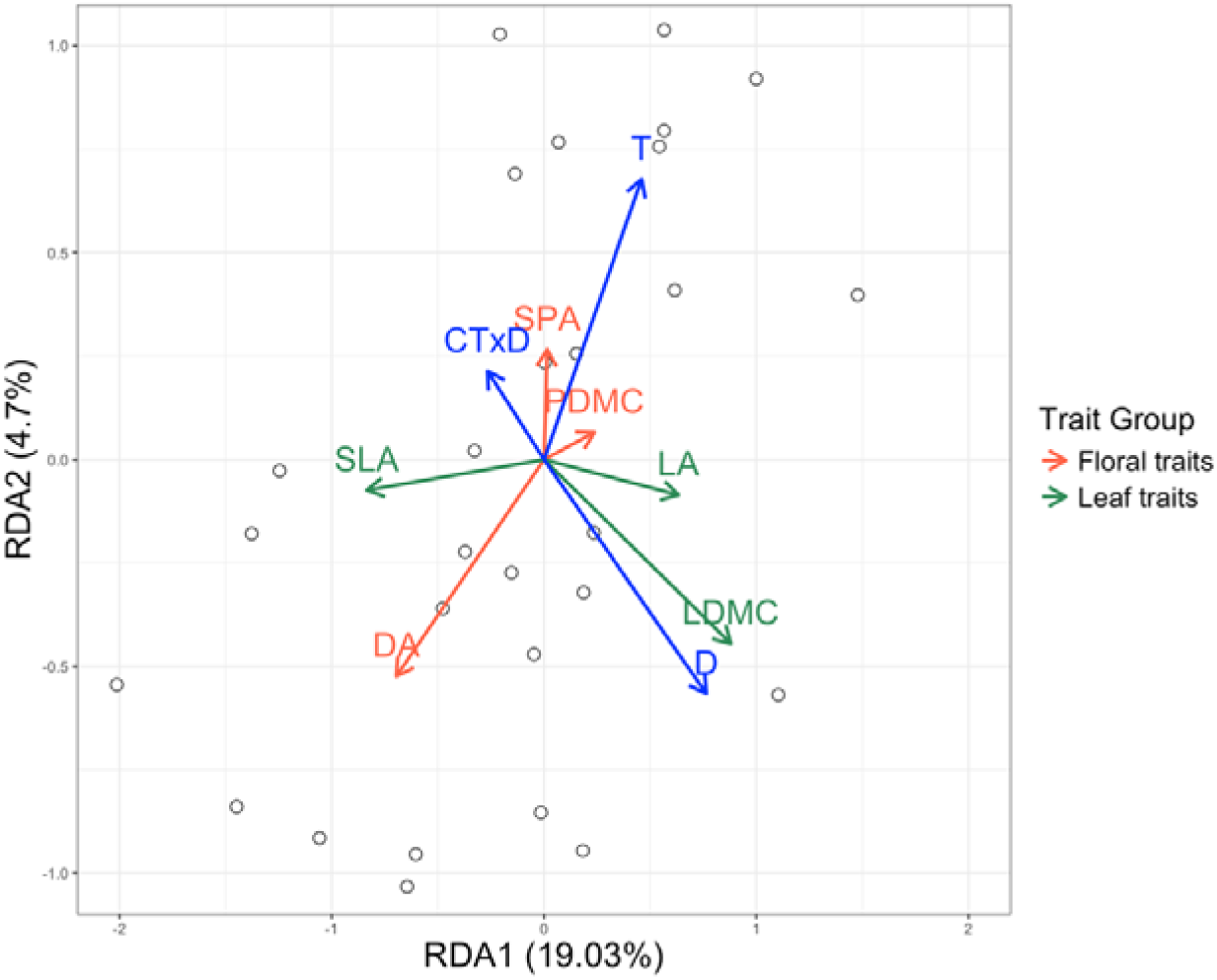
RDA plot of *L. corniculatus*. Significant climate change factors have been shown in blue while plant traits have been coloured based their group (as shown in legend). ; DA, Display Area; SPA, Specific Petal Area; PDMC, Petal Dry Matter Content; LA, Leaf Area; SLA, Specific Leaf Area; LDMC, Leaf Dry Matter Content; T, Warming (+3°C); D, Drought; CT×D, Elevated CO_2_, warming and drought.

Drought and temperature were the primary drivers of variation along the first RDA axis, which accounted for 19.03% of the variation in trait responses. Display Area (DA) and leaf traits (Leaf Area (LA), Specific Leaf Area (SLA), and Leaf Dry Matter Content (LDMC)) varied along this gradient, with increasing drought being associated with an increase in Leaf Area and Leaf Dry Matter Content (LDMC) alongside a decrease in Specific Leaf Area (SLA). Temperature was diametrically opposite to Display Area (DA), indicating that increasing temperature levels would lead to a reduction in Display Area (DA). The second RDA axis accounted for 4.7% of trait variation and was related to CT×D. Display Area (DA) was well-represented on this axis, indicating that the most extreme condition, CTD, would lead to a decrease in Display Area (DA).

The univariate analyses of *L. corniculatus* traits revealed that drought had the most consistent and significant effects (Fig. 2), increasing Leaf Area (LA), Leaf Dry Matter Content (LDMC) while decreasing Specific Leaf Area (SLA) (Fig. 3d, 3e and 3f). CO_2_ induced variation in two out of the three leaf traits, reducing Specific Leaf Area (SLA) and increasing Leaf Dry Matter Content (LDMC) (Fig. 3a and 3b). The temperature effect was more trait specific as it had a significant effect only on Display Area (DA) (Fig. 3c), causing a substantial decrease in Display Area (DA) under higher temperature. In terms of interactions, C×T overall caused a decrease in Display Area (DA), as warming negated and reversed the positive effect of CO_2_ (Fig. 3g). In the case of Specific Leaf Area (SLA), the decrease caused by CO_2_ became stagnant with increasing temperature (Fig. 3h). In the most extreme treatment (CT×D), drought had a buffering effect on increase of Leaf Dry Matter Content (LDMC) by CO_2_ and temperature (Fig. 3i).

**Fig. 2.**
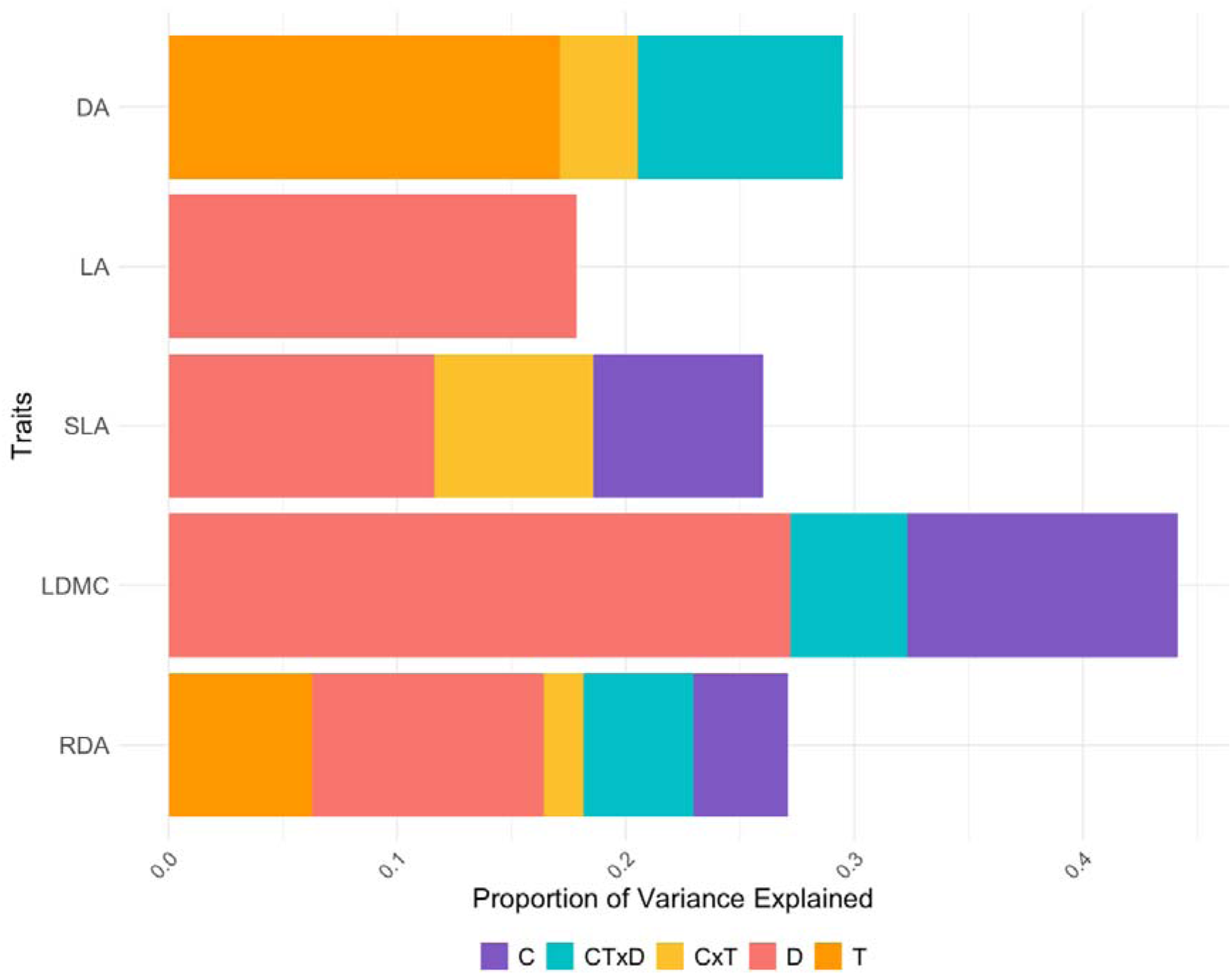
Variance partitioning plot for *L. corniculatus* showing magnitude of variation explained by the different climate change factors. C, Elevated CO_2_ (+300ppm); T, Warming (+3°C); D, Drought; C×T, Elevated CO_2_ combined with warming; CT×D – Elevated CO_2_, warming and drought; DA, Display Area; LA, Leaf Area; SLA, Specific Leaf Area; LDMC, Leaf Dry Matter Content. RDA results are of multivariate analysis. Only the significant variables of p < 0.05 are shown. Specific Petal Area and Petal Dry Matter Content do not have any significant predictors and are thus not shown.

**Fig. 3.**
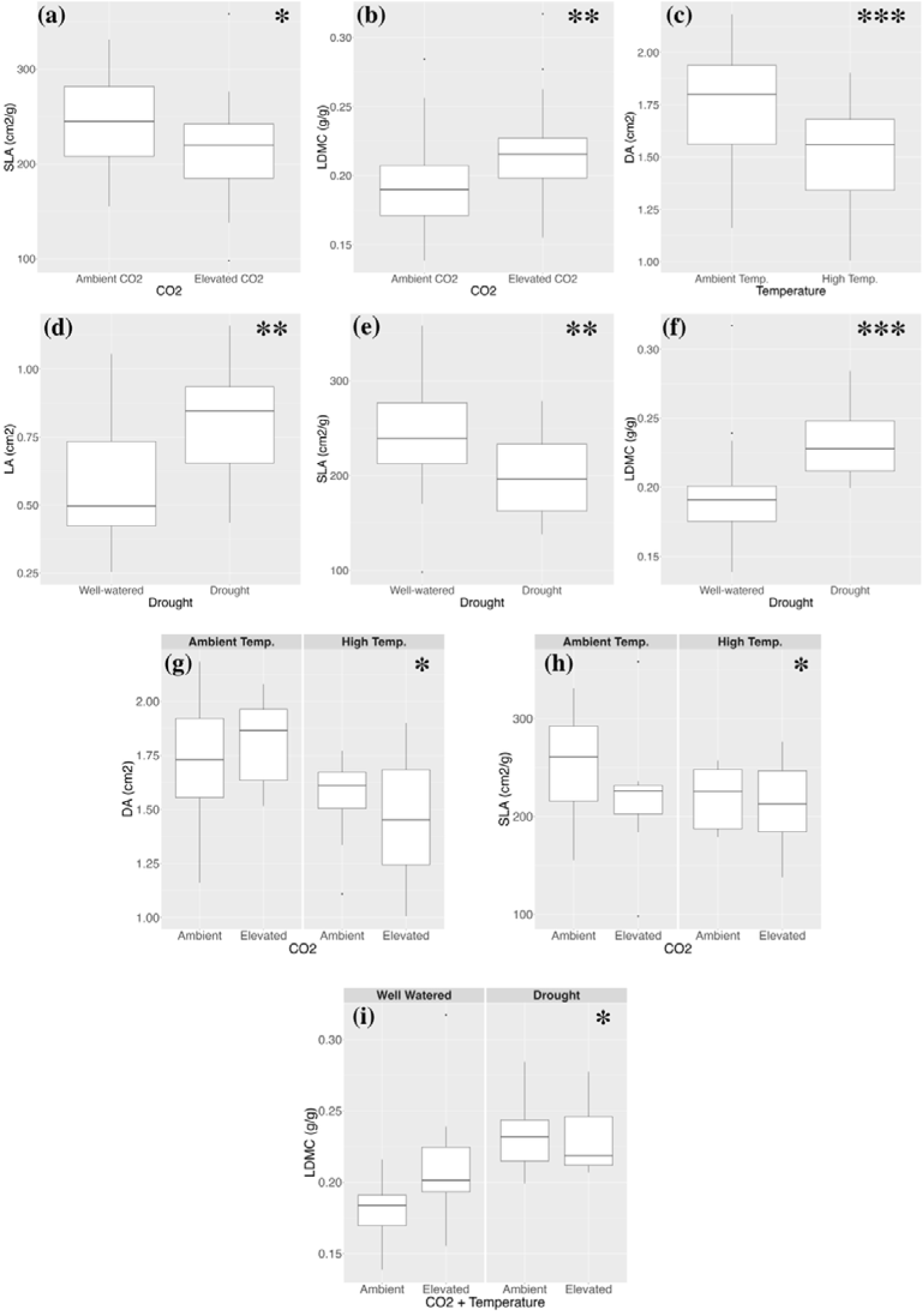
Boxplots illustrating the individual and combined effects of climate change factors on plant traits of *L. corniculatus*. Only the significant effects are shown with the significance level of the effect of the plotted factor or their interaction indicated in the top right corner (* = 0.05; ** = 0.01; *** = < 0.001). Functional trait studied denoted in the Y-axis while treatment is denoted in the X-axis. For legend of plant traits see Fig. 2.

### Crepis capillaris

The RDA analysis of *C. capillaris* showed that the climate change factors accounted for 20% of the total trait variation (p = 0.003 for the overall model) and drought (7.7%) and C×T (4.5%) having significant effect on the plant traits (Fig. 4).

**Fig. 4.**
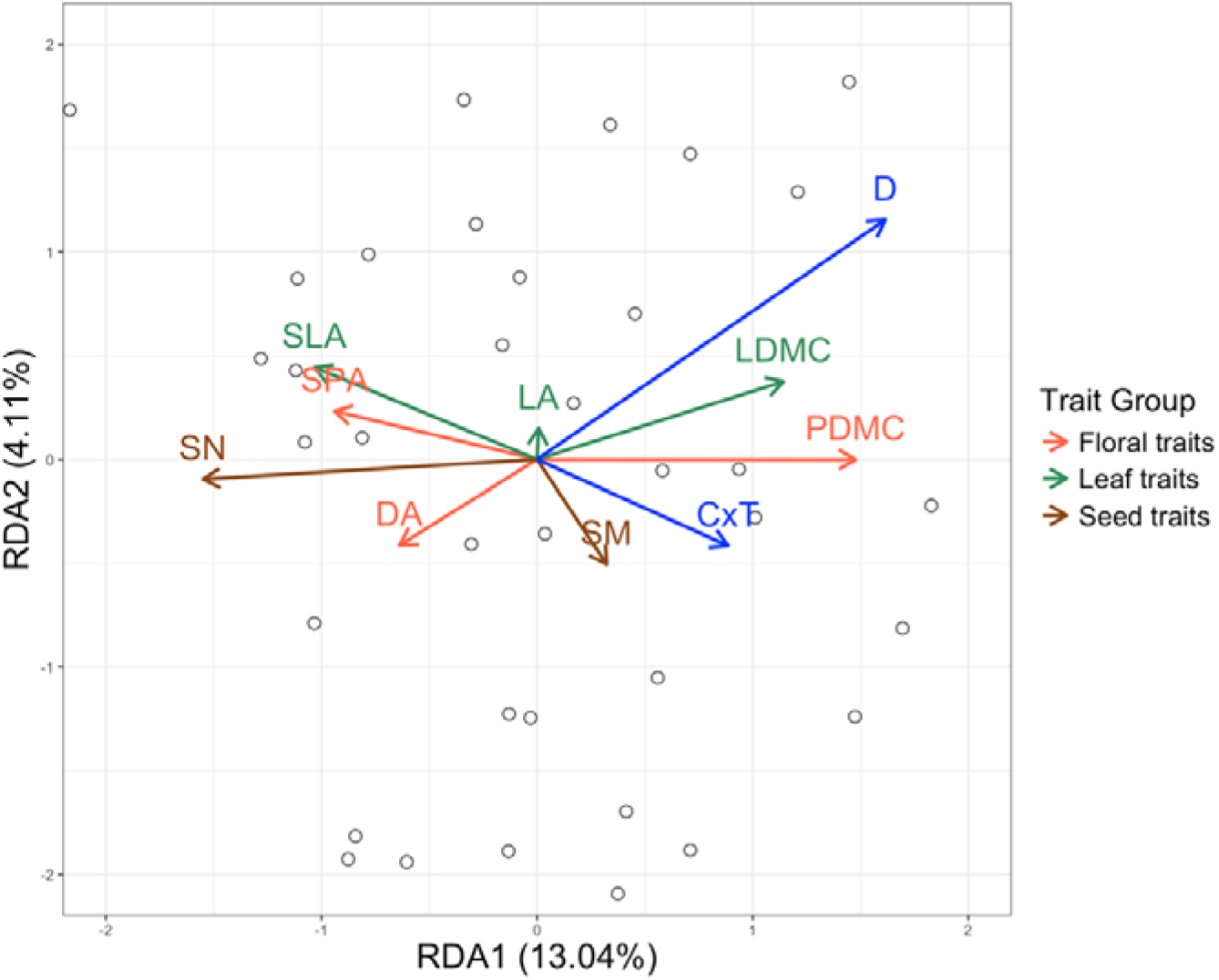
RDA plot of *C. capillaris*. Significant climate change factors have been shown in blue while plant traits have been coloured based their group (as shown in legend). DA, Display Area; SPA, Specific Petal Area; PDMC, Petal Dry Matter Content; LA, Leaf Area; SLA, Specific Leaf Area; LDMC, Leaf Dry Matter Content; SN, Seed Number; SM, Seed Mass; D, Drought; C×T, Elevated CO_2_ combined with warming.

These two climate change factors were highly correlated in the first RDA axis (13%), with drought being the primary and C×T the secondary contributor to variation along this axis. The constrained variation in floral traits (Display Area (DA), Specific Petal Area (SPA), and Petal Dry Matter Content (PDMC)), leaf traits (Specific Leaf Area (SLA) and Leaf Dry Matter Content (LDMC)), and Seed Number (SN) was chiefly represented along this axis. The patterns indicated that drought had a positive relationship with the Dry Matter Content traits (Leaf Dry Matter Content (LDMC) and Petal Dry Matter Content (PDMC)) while having a negative relationship with all other traits. Among these traits, Seed Number (SN) showed the strongest inverse relationship with drought. The second RDA axis accounted for 4% of the total variation, with temperature as the dominant factor. Leaf Area (LA) was best represented along this axis, possibly indicating that increasing temperature would cause a decrease in Leaf Area (LA), although this relationship was not significant.

The findings in the univariate analysis reflected the general patterns observed in the RDA. Similar to *L. corniculatus*, trait variation in *C. capillaris* was primarily dictated by drought, which was the only climatic variable having a significant effect on the majority of plant traits. Its magnitude of variation was highest in Seed Number (SN), followed by Petal Dry Matter Content (PDMC) and then the other traits (Specific Petal Area (SPA), Leaf Dry Matter Content (LDMC), and Display Area (DA)) (Fig. 5). An increase in drought levels caused a significant decrease in Display Area (DA), Specific Petal Area (SPA), and Seed Number (SN) while leading to an increase in Petal Dry Matter Content (PDMC) and Leaf Dry Matter Content (LDMC) (Fig. 6a – 6e).

**Fig. 5.**
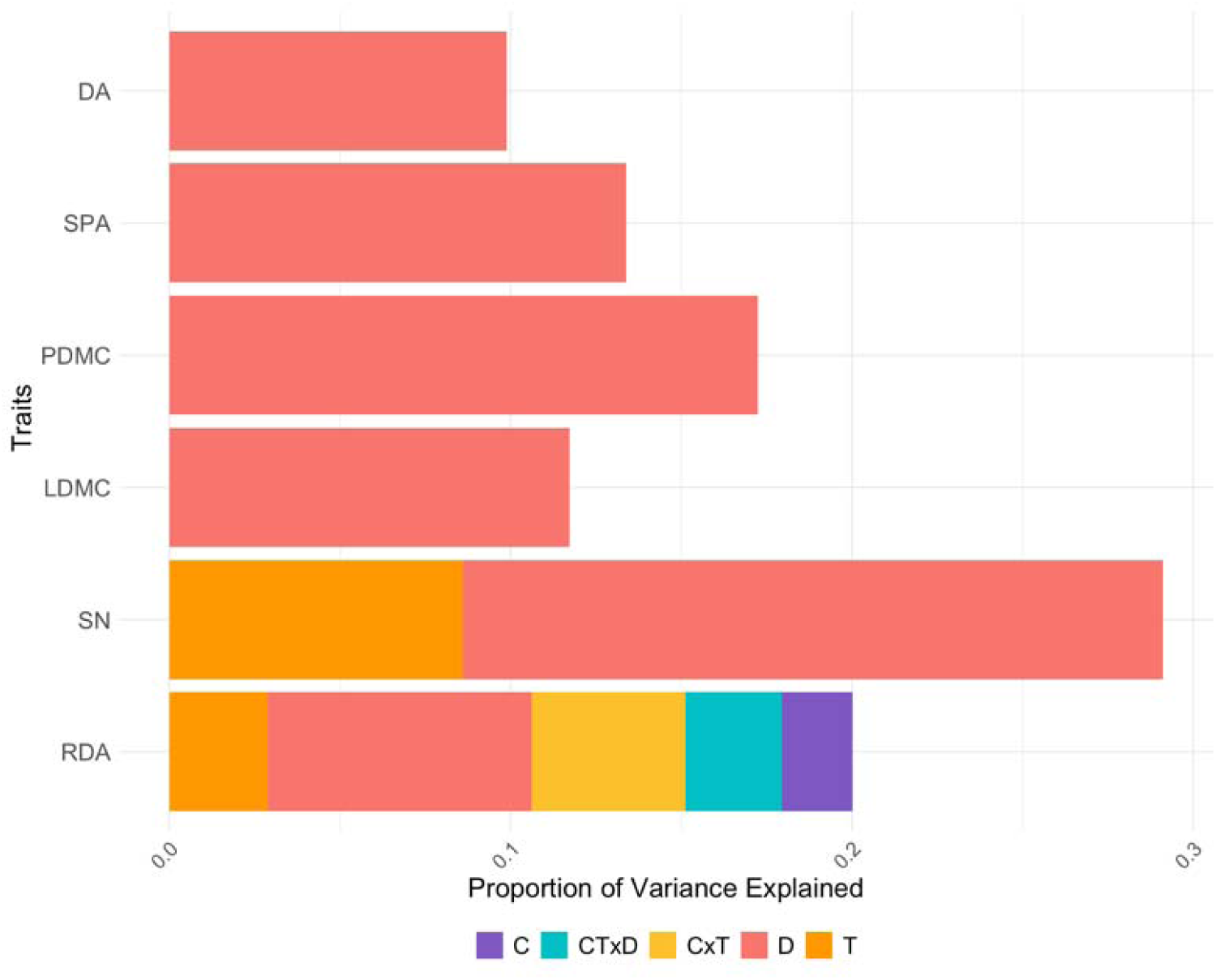
Variance partitioning plot for *C. capillaris* explaining magnitude of variation in relation to the three climate change factors and their interactions. C, Elevated CO_2_ (+300ppm); T, Warming (+3°C); D, Drought; C×T, Elevated CO_2_ combined with warming; CT×D – Elevated CO_2_, warming and drought; DA, Display Area; SPA, Specific Petal Area; PDMC, Petal Dry Matter Content; LDMC, Leaf Dry Matter Content; SN, Seed Number. RDA results are of multivariate analysis. Only the significant variables of p < 0.05 are shown. Specific Leaf Area does not have significant predictors.

**Fig. 6.**
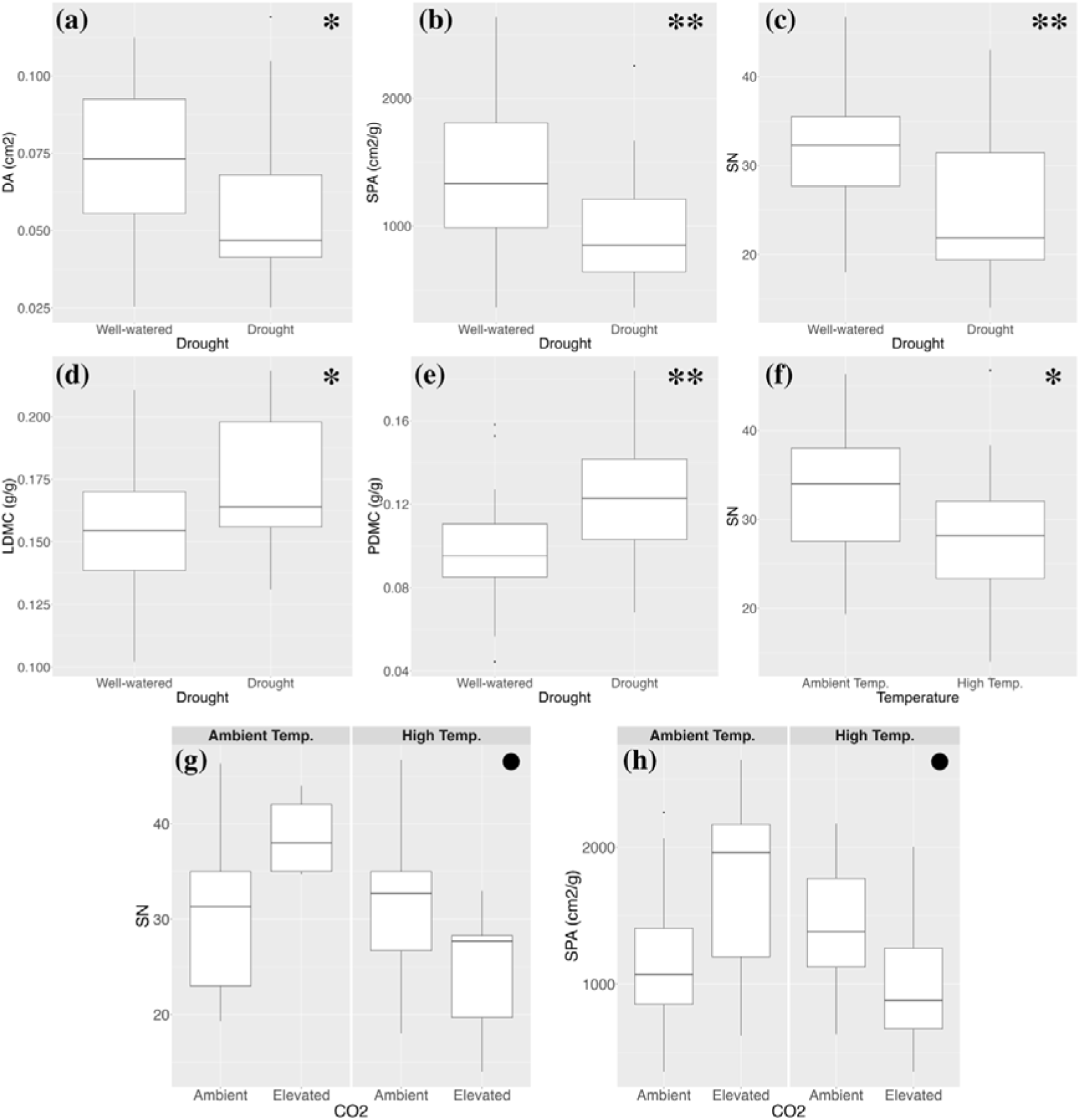
Boxplots illustrating the effects of climate change factors on plant traits of *C. capillaris*. Only the significant effects have been shown with the significance level of the effect of the plotted factor or their interaction is indicated in the top right corner (· = <0.1; * = 0.05; ** = 0.01). Functional trait studied denoted in the Y-axis while treatment is denoted in the X-axis. For legend of plant traits see Fig. 5.

Similar to *L. corniculatus*, effect of temperature in *C. capillaris* is also trait-specific and caused a significant effect only for Seed Number (SN), which was negatively related to warming (Fig. 6f). In terms of interactions, although under ambient temperature conditions, increase in CO_2_ intensity causes increase in Specific Petal Area (SPA), warming is shown to negate the positive effect of CO_2_ with CT showing slight reduction in Specific Petal Area (SPA) despite the positive effect of CO_2_ (Fig. 6g). This pattern is mirrored in the effect of CO_2_ and temperature on Seed Number (SN) (Fig. 6h).

### Covariation Between Plant Traits

The initial Principal Component Analysis (PCA) of both species revealed two independent axes of variation among leaf and floral traits. In *L. corniculatus*, the first two axes accounted for 64% of the variation (Fig. 7a). The first axis accounted for 35.2% of the variation and was primarily associated with leaf traits (Specific Leaf Area (SLA) and Leaf Dry Matter Content (LDMC)). The second axis accounted for 28.8% of the variation and was linked to floral traits (Specific Petal Area (SPA) and Petal Dry Matter Content (PDMC)). Specific area traits (Specific Leaf Area (SLA) and Specific Petal Area (SPA)) were negatively correlated with their dry matter counterparts (Leaf Dry Matter Content (LDMC) and Petal Dry Matter Content (PDMC)). Regarding floral traits, Display Area (DA) occurred closely with Specific Petal Area (SPA), and together they were orthogonal to the leaf traits, where Leaf Area (LA) occurred closely with Leaf Dry Matter Content (LDMC).

**Fig. 7.**
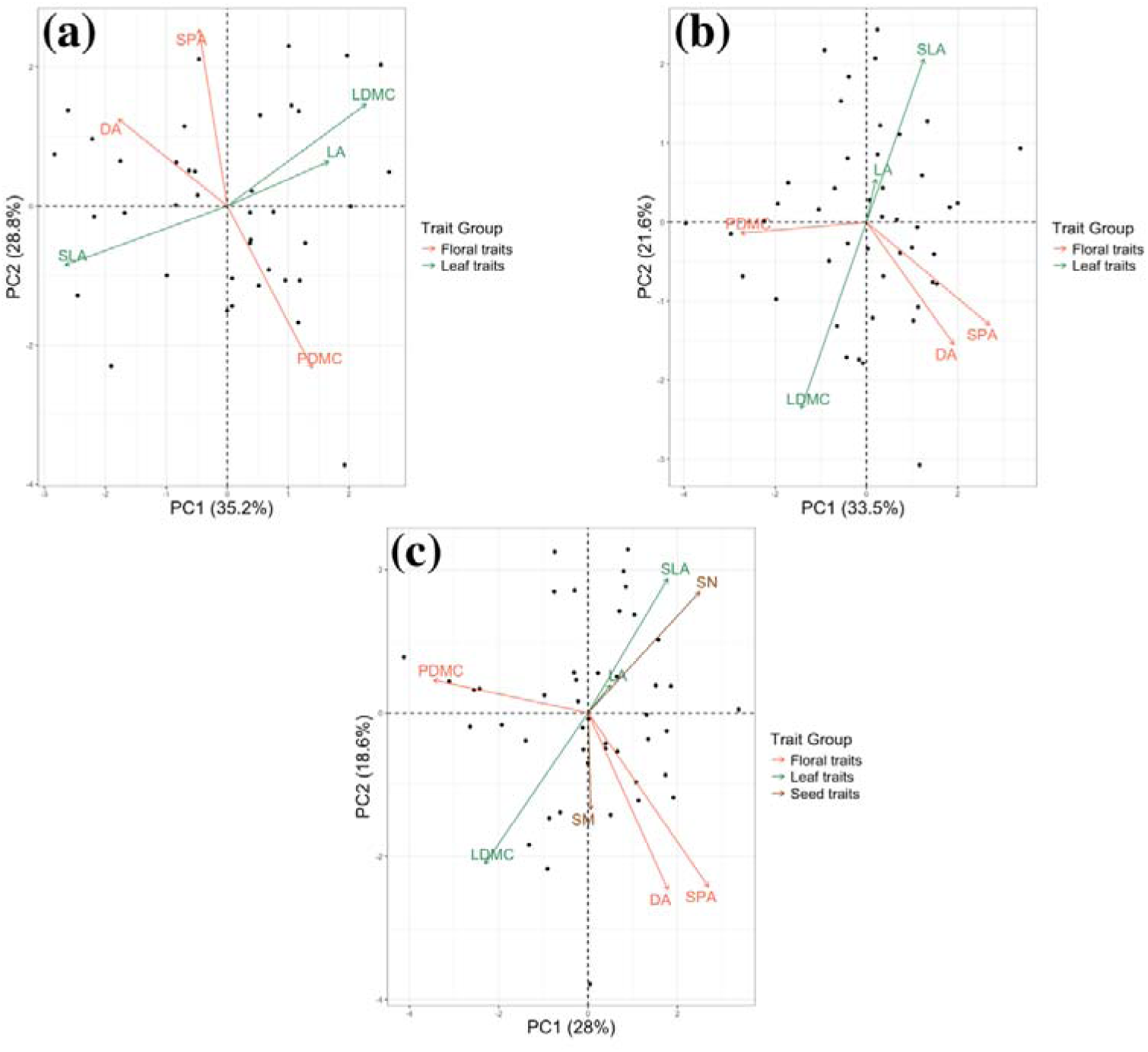
PCA plot of (a) *L. corniculatus* (b) *C. capillaris* (without seed traits) and (c) *C. capillaris* (with seed traits). DA, Display Area; SPA, Specific Petal Area; PDMC, Petal Dry Matter Content; LA, Leaf Area; SLA, Specific Leaf Area; LDMC, Leaf Dry Matter Content; SN, Seed Number; SM, Seed Mass.

In *C. capillaris*, the first two principal components accounted for 55.2% of the total trait variation (when analyzed without seed traits) (Fig. 7b). The first axis was dominated by floral traits (Display Area (DA), Specific Petal Area (SPA), and Petal Dry Matter Content (PDMC)) (33.6%), while the second axis was dominated by leaf traits (Leaf Area (LA), Specific Leaf Area (SLA), and Leaf Dry Matter Content (LDMC)). Similar to Display Area (DA) in *L. corniculatus*, Display Area (DA) in *C. capillaris* occurred closely with Specific Petal Area (SPA) and was orthogonal to leaf traits (Specific Leaf Area (SLA) and Leaf Dry Matter Content (LDMC)). Specific Leaf Area (SLA) and Leaf Dry Matter Content (LDMC) were also negatively correlated.

When seed traits (Seed Number (SN) and Seed Mass (SM)) were added to the PCA (Fig. 7c), the overall patterns of dominance remained the same. Seed Number (SN) was negatively correlated with the Dry Matter Content traits (Petal Dry Matter Content (PDMC) and Leaf Dry Matter Content (LDMC)). The loadings for Seed Mass (SM) consistently had low absolute values across all principal components, with a maximum of -0.3 on PC3. This implied that Seed Mass (SM) played a negligible role in defining the principal components.

Network analysis of the two species confirms the findings from the PCAs while offering some additional insights by revealing the strength and structure of trait interrelationships. In *L. corniculatus*, the most connected and most central trait was Petal Dry Matter Content (PDMC) (Fig. 8a) which connected the other floral traits, Specific Petal Area (SPA) and Display Area (DA) while having a stronger connection with the former (Supplementary Information Table S3). Both floral and leaf traits independently clustered with the leaf traits having stronger interactions among each other than the floral traits (Fig. 8). These findings contrast with that of *C. capillaris* (Fig. 8b) where dry matter content traits, specifically Leaf and Petal Dry Matter Content form a cluster alongside Specific Leaf Area (SLA) and Seed Number (SN). Specific Petal Area (SPA) and Petal Dry Matter Content (PDMC) dominate the network both in degree and betweenness. Traits such as Leaf Area (LA) and Seed Mass (SM) do not play any role in the network.

**Fig 8.**
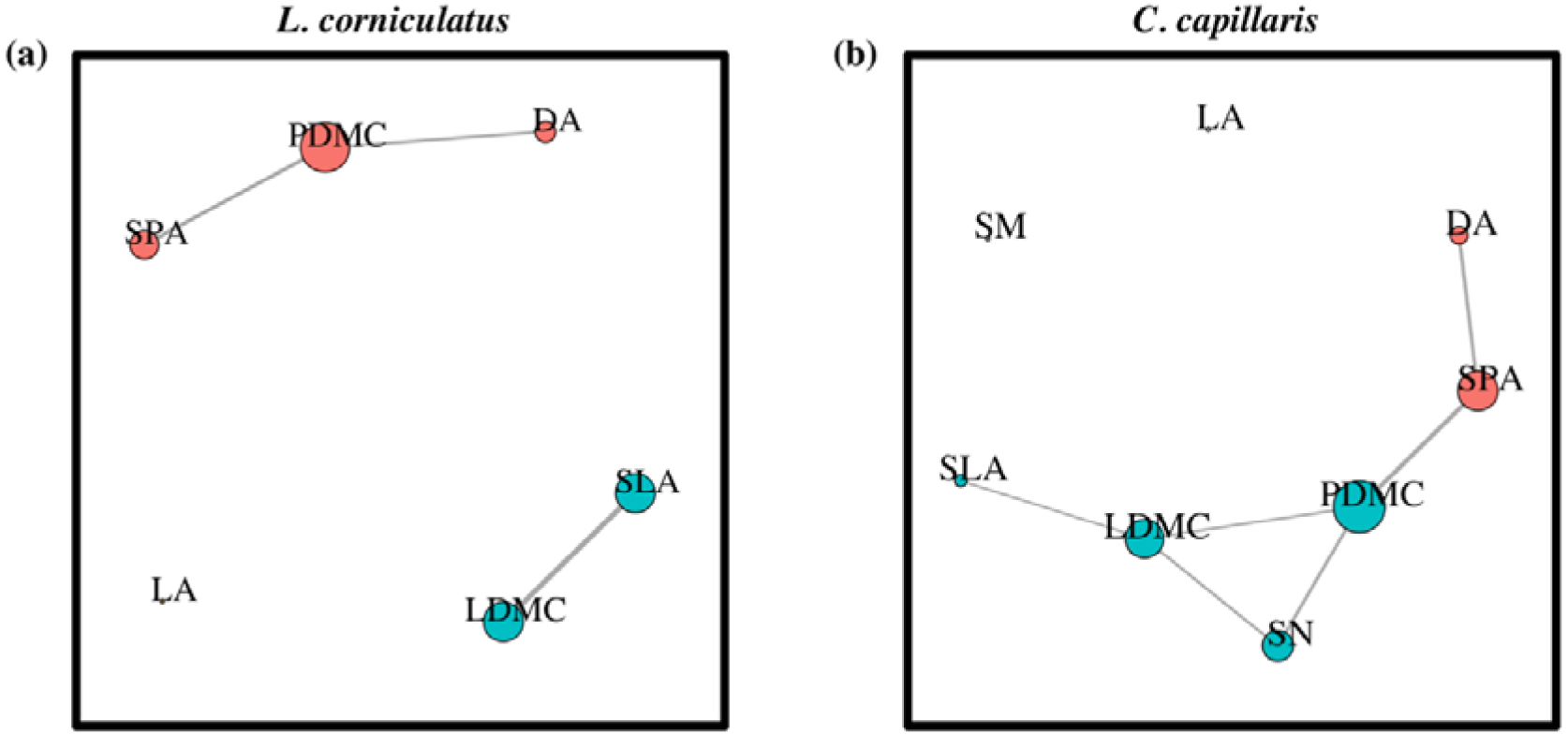
Trait correlation network for (a) *L. corniculatus* and (b) *C. capillaris*. Node colours indicate separate clusters or “modules” while node size shows betweenness. Correlation strength is shown by line thickness and distance between traits. DA, Display Area; SPA, Specific Petal Area; PDMC, Petal Dry Matter Content; LA, Leaf Area; SLA, Specific Leaf Area; LDMC, Leaf Dry Matter Content; SN, Seed Number; SM, Seed Mass.

### Coordinated Changes in Traits in Response to Climate change Factors

When analysing the covariation of plant traits under each climatic regime separately, we found that different treatments led to coordinated adjustments in traits. However, the coordinated responses were not similar across species. In *L. corniculatus*, the most extreme treatment (C1T1D1) caused a major coordinated adjustment in traits (Fig. 9a), with PC1 being the only significant axis explaining 65.3% of the total variation. Variation in this axis was dictated by Specific Petal Area (SPA) and Leaf Dry Matter Content (LDMC). Traits in the other treatments did not show any significant covariations.

**Fig. 9.**
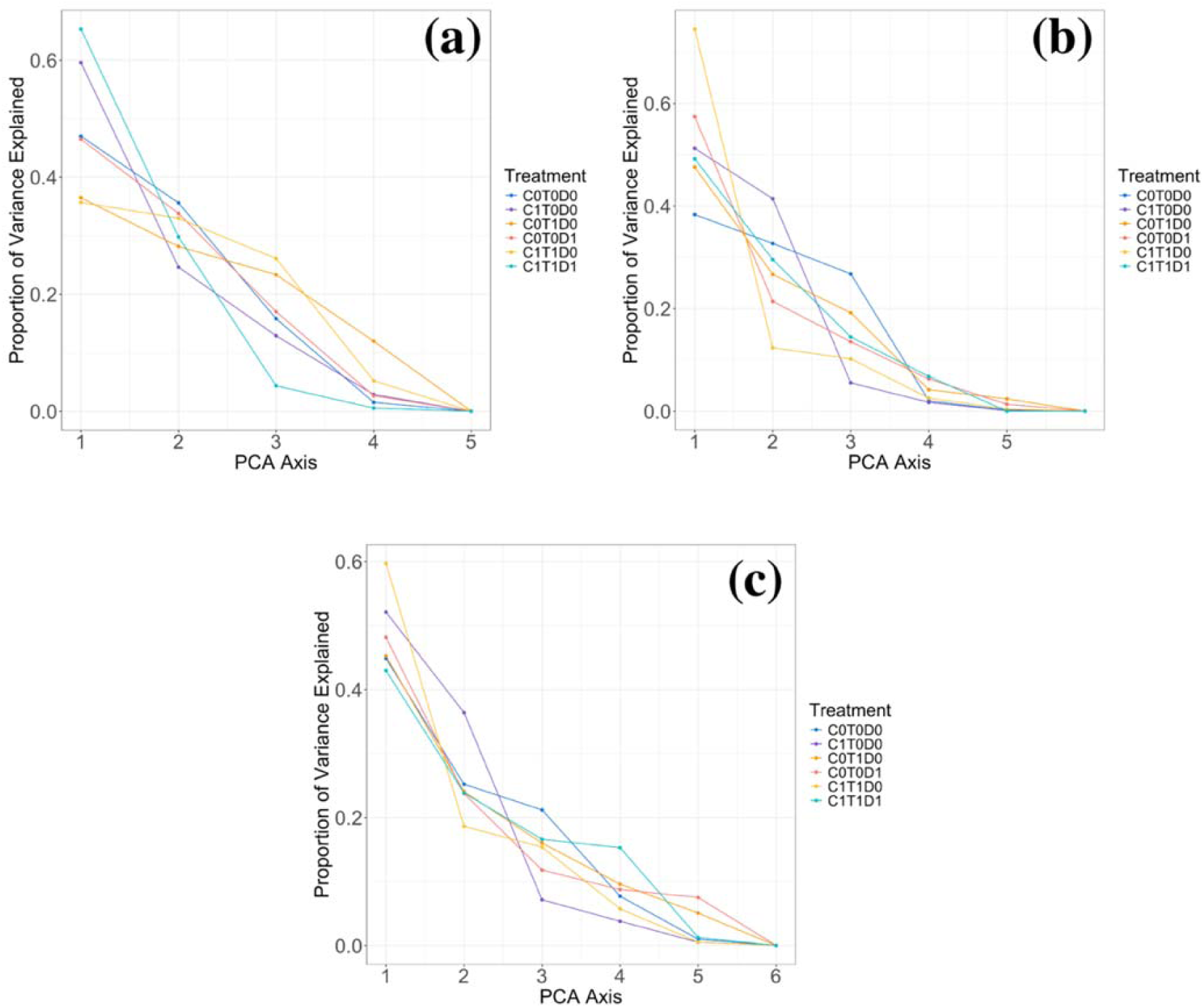
Variation explained in the traits by each of the PCA axes under different treatments in (a) *L. corniculatus*, (b) *C. capillaris* (without Seed Traits) and (c) *C. capillaris* (with Seed Traits). C0T0D0 - Ambient Control, C1T0D0 – Elevated CO_2_ (+300ppm), C0T1D0 – Warming (+3°C), C0T0D1 – Drought, C1T1D0 – Elevated CO_2_ combined with warming, C1T1D1 – Elevated CO_2_, warming and drought.

These patterns were partially mirrored in *C. capillaris* (Fig. 9b). Under the combined elevated CO_2_ and warming treatment, PC1 accounted for 74.5% of the total trait variance and was statistically significant. Specific Petal Area (SPA), Petal Dry Matter Content (PDMC), Leaf Area (LA) and Specific Leaf Area (SLA) all have strong loadings on PC1, with SPA, LA and SLA also showing significant correlations with the axis. In contrast, under the elevated CO_2_ treatment alone, only PC2 that was significant, explaining 41.4% of the variance. In this case, Leaf Area (LA) and Leaf Dry Matter Content (LDMC) loaded significantly on PC2, with LDMC also exhibiting a significant correlation with the axis. No significant trait covariation was observed in the remaining treatment combinations. These PCA analyses should be interpreted with caution as the small sample size means that the number of variables analysed was equal to or greater than the number of observations. This increases the risk of overfitting and reduces the robustness of the PCA outcomes. To mitigate this limitation, we interpret only on the treatments which consistently showed statistical significance across multiple randomizations. This strategy helped ensure that the observed patterns of trait covariation were not artefacts of sampling variability, but reflected biologically meaningful responses-albeit with caution to their generalizability.

## Discussion

The results of this study show that climate change simulated for multiple years significantly impacts not only vegetative traits but also reproductive traits such as floral and seed traits although responses vary between species. Our study supported (H1) indicating that plant traits respond more to drought more consistently than to elevated CO_2_ and warming. Contrary to (H2), the negative effects of temperature were not mitigated by elevated CO_2_ conditions. Finally, in line with (H3) climate change factors and their interactions exerted a strong filtering on covariation of plant traits, albeit this phenomenon was species-specific and our data have limited strength to prove this unequivocally.

### Response of Plant Functional Traits to Individual Climate change Factors

Climate change factors significantly influenced the functional traits of both *L. corniculatus* and *C. capillaris*, though the responses varied across traits and species. In *L. corniculatus*, drought was the dominant factor, but other climate change factors also affected two of the three leaf traits (Specific Leaf Area and Leaf Dry Matter Content) and one floral trait (Display Area). In *C. capillaris*, drought played a major role, significantly impacting all floral traits (Display Area, Specific Petal Area, and Petal Dry Matter Content), as well as one leaf trait (Leaf Dry Matter Content) and one seed trait (Seed Number). Temperature was the only other significant factor, affecting only Seed Number. Species-specific responses have been reported in many previous studies (Ainsworth and Long, 2005; Descamps *et al*., 2020; Kuppler and Kotowska, 2021; Bernauer *et al*., 2024).

The finding that drought is the dominant factor affecting economic traits of both species aligns with our first hypothesis (H1). Drought stress has been consistently shown to reduce traits such as Leaf Area, Specific Leaf Area, Flower size, and Seed Number (Lidon, 2012; Wellstein *et al*., 2017; Matesanz *et al*., 2020; Descamps *et al*., 2021; Höfer *et al*., 2023), while increasing Leaf Dry Matter Content (Jung *et al*., 2014). It is reflecting a shift toward more carbon-demanding structures with higher construction costs per unit area. A notable exception in our study is the response of Leaf Area. In *L. corniculatus*, Leaf Area increased under drought, contrary to what was found by most prior studies (e.g., Westoby *et al*., 2002; McDonald *et al*., 2003; Yates *et al*., 2010; Bjorkman *et al*., 2018; Zhou *et al*., 2020). Our finding is supported by a study of *Limonium* species, where larger leaves despite being considered more carbon-demanding in terms of construction cost were associated with higher growth capacity and higher water use efficiency (Conesa, Mus and Galmés, 2019). Our study demonstrated a more conservative strategy involving higher carbon investment, in which plants increased dry matter content traits (such as Leaf Dry Matter Content and Petal Dry Matter Content) while reducing tissue construction cost-related traits (such as Specific Leaf Area and Specific Petal Area). This shift suggests a higher investment in structural tissues, aligning with the plant’s adaptive response to environmental stress (Poorter *et al*., 2009). These tissues, although more costly to produce, are more durable and enhance plant resistance under environmental stress (Gong, Cui and Gao, 2020; Wang *et al*., 2022; Akram *et al*., 2023). Smaller plant organs generally require less water to maintain turgor pressure, which can improve plant water status during drought stress by reducing overall transpiration and hydraulic demand (Galen, Sherry and Carroll, 1999; CHAVES *et al*., 2002). Although this strategy is cost-effective for temporary survival, the reduction in flower size, a crucial visual cue for pollinators, could potentially decrease pollinator visitation rates and thereby reducing population growth rates (Descamps, Quinet and Jacquemart, 2021). However, a decrease in Specific Petal Area may also suggest increased floral longevity due to reduced flower maintenance costs (Spigler and Woodard, 2019). This could mitigate the potential negative effects of smaller flower size. While no general patterns of decreasing visitation rates by pollinators under water deficit conditions have been identified (reviewed in Kuppler and Kotowska, 2021), the potential impact of floral trait variation on pollinator behaviour requires further attention.

Warming influenced only two plant traits: it reduced floral display area in *L. corniculatus* and seed number in *C. capillaris*, without affecting any resource economic traits. Although there are many previous studies which predicted a significant linear relationship between leaf traits (Leaf Area, Specific Leaf Area and Leaf Dry Matter Content) and temperature (e.g., Atkin *et al*., 2006; Poorter *et al*., 2010; Fontana *et al*., 2017; Dostálek, Rokaya and Münzbergová, 2020), our findings on leaf traits were more in line with a study conducted in the Bavarian Alps consisting of 223 species (Rosbakh, Römermann and Poschlod, 2015) where a significant relationship between Specific Leaf Area and temperature was found only at the community level with no response at either the species or population level. This could be because of the predominant effect of drought on vegetative traits as water availability is a greater limiting factor than temperature in temperate climates (Fahad *et al*., 2017; Zhou *et al*., 2017). In regard to floral traits, although warming did not alter the per area structural investment in either species, it led to reduced display area in one of them. This response may be attributed to the greater sensitivity of reproductive development to abiotic stresses as compared to vegetative growth (Zinn, Tunc-Ozdemir and Harper, 2010; Snider and Oosterhuis, 2011). Under warming conditions, this heightened sensitivity can lead to reductions in floral organ size, potentially through altered expression of genes in floral morphogenesis (Zinn, Tunc-Ozdemir and Harper, 2010; Smith and Zhao, 2016). In light of this, species capable of maintaining and increasing floral display size under elevated temperatures have a greater probability of surviving in future climate scenarios over species which cannot. However, this strategy may involve trade-offs between display size and floral longevity, as larger flowers typically require greater construction costs and may have shorter lifespans (Reich, 2014). Plants that can flexibly adjust their display size and flower longevity can enhance pollinator attraction when pollinators are rare by increasing their display size and limit self-pollination when pollinators are common by reducing their display size (Harder and Johnson, 2005).These dynamics highlight the importance of floral trait plasticity-not just in size but also in timing and lifespan-as a potential adaptive response to climate change. While our study did not include measurements of flower longevity, incorporating this trait in future research would provide valuable insights into how plants balance reproductive investment under environmental stress.

In our experiment, elevated CO_2_ levels resulted in a decrease in Specific Leaf Area (SLA) and increase in Leaf Dry Matter Content in only one species (*L. corniculatus*). Similar to the drought response, this pattern suggests a shift toward a more carbon-demanding and conservative strategy. While several studies confirm our findings of decrease in Specific Leaf Area and increase in Leaf Dry Matter Content under elevated CO_2_ (e.g., Centritto, Lee and Jarvis, 1999; Stiling and Cornelissen, 2007; Robinson, Ryan and Newman, 2012; Poorter *et al*., 2022), others partially contradict them, showing increased Specific Leaf Area (SLA) under elevated CO_2_ (Duan *et al*., 2018; Temme *et al*., 2019). This variation may be due to differences in resource-use strategies among plants with different growth forms. Perennials like *L. corniculatus* often have thicker leaves with lower Specific Leaf Area (SLA) and higher Leaf Dry Matter Content, leading to slower growth rates and lower resource acquisition compared to faster-growing annuals (Poorter *et al*., 2009; Reich, 2014). When exposed to favourable conditions (such as elevated CO_2_), perennials may not exhibit the same growth responses as annuals because their strategies prioritize survival and maintenance over rapid growth. In our study, elevated CO_2_ level had no effect on floral traits. This is in line with (Bernauer *et al*., 2024) while other studies have reported both increase and decrease of flower size under elevated CO_2_ (Niu *et al*., 2000; Hoover *et al*., 2012; Maia *et al*., 2023). Overall, our findings highlight how environmental stressors such as drought and elevated CO- drive shifts in plant economic strategies. Traits like Specific Leaf Area and Specific Petal Area serve as indicators of tissue construction cost, with lower values reflecting more carbon-demanding and more structurally robust tissues. Conversely, higher dry matter content traits (LDMC and PDMC) signal a conservative strategy prioritizing durability over rapid resource acquisition. These shifts underscore the importance of integrating plant economics frameworks when interpreting trait responses to climate change.

### Interactive Effects of Climate change Factors on Plant Traits

Regarding the combined effects of climate change factors on plant traits, we hypothesized (H2) that the negative effects of either warming or drought would be partially mitigated by elevated CO_2_. This hypothesis was not supported by our observations. The combination of elevated CO_2_ and warming reduced Display Area of *L. corniculatus*, and on Specific Petal Area and Seed Number in *C. capillaris*. Similar findings have been reported in other studies. For instance, Li et al. (2019), in their study on *Morus alba* L., found that elevated CO- and temperature together reduced the number and biomass of male inflorescences, while positively affecting female inflorescences. Notably, the fresh weight of male inflorescences under combined elevated CO- and warming was lower than under warming alone, mirroring the pattern observed in our study (Li *et al*., 2019). Other research has also shown that elevated CO- does not offset the negative effects of warming on seed yield, likely due to reduced photosynthetic efficiency under heat stress, which can lead to ovule abortion and impaired reproductive development (Ruiz-Vera *et al*., 2015; Notarnicola *et al*., 2023).

The combination of elevated CO_2_, warming and drought (CT×D) significantly affected only one economic trait in only one species, i.e. Leaf Dry Matter Content (LDMC) in *L. corniculatus*. While plants exposed to elevated CO- and temperature exhibited higher Leaf Dry Matter Content than those in ambient conditions, drought-stressed plants showed a significantly greater Leaf Dry Matter Content than both ambient and elevated CO- + temperature. Increased Leaf Dry Matter Content might be attributed to improved water-use efficiency, driven by reduced Specific Leaf Area (Wellstein *et al*., 2017) and increased investment in non-structural carbohydrates (Blumenthal *et al*., 2020; Rodríguez-Alarcón, Tamme and Carmona, 2022). Changes in palisade parenchyma cell composition— specifically increases in cell number and size under drought—may also contribute to reduced Specific Leaf Area and increased Leaf Dry Matter Content (Peña-Rojas *et al*., 2024), reinforcing the link between anatomical adaptation and economic trait shifts.

### Trait Covariation Patterns and Effect of Climate change Factors on Covariations

Our findings on trait covariation patterns reveal that floral traits of *L. corniculatus* are largely independent of leaf traits and this is in-line with (E-Vojtkó *et al*., 2022). In *C. capillaris* however, there seems to be an interesting link between Petal and Leaf Dry Matter Content with both traits also having Seed Number as a mediator. Although previous studies do not specifically use these two traits, overall plant dry matter has been found to be positively correlated with seed number (Vanderlip *et al*., 2004; Aparna *et al*., 2017). This would be due to the higher dry matter accumulation enabling plants to store and mobilize more resources for seed development, leading to increased seed yield (Liu *et al*., 2020). In regard to the effects of climate change factors on trait covariation, we hypothesized (H3) that environmental extremes exert a strong coordinated response in plant functional traits. This phenomenon was found to be true for only one of our two species. This differential coordinated response of plant functional traits may reveal interesting insights into the adaptability and distribution of plant species across different habitats.

Functional traits of *L. corniculatus* showed the most coordinated response in the most extreme conditions (C1T1D1). This is in-line with previous studies (Lozano *et al*., 2020; Mueller, Kray and Blumenthal, 2024). Drier, warmer habitats have been found to exert stronger selection pressures on plant traits to enhance water-use efficiency, heat tolerance and resource conservation which eventually leads to a stronger selection for trait coordination (Blumenthal *et al*., 2020; Mueller, Kray and Blumenthal, 2024). *C. capillaris* showed coordinated trait responses primarily under elevated CO- alone (C1T0D0) and in combination with elevated temperature (C1T1D0). The absence of a coordinated trait response to drought in *C. capillaris*, unlike in *L. corniculatus*, may contribute to the differences in their ecological distribution and adaptive capacity. *L. corniculatus* is widely distributed across diverse climatic zones, thriving in temperate, tropical and sub-tropical climates while also occurring in certain regions of the polar and sub-polar zones. Although *C. capillaris* is also broadly distributed, it is primarily restricted to the temperate regions of Europe and North America while having some coverage in the sub-tropical zones of Africa and Australia (“Plants of the world online,” 2024). These findings suggest that trait coordination under stress may support ecological adaptability and range expansion. However, it is unlikely to be the sole factor determining species distribution, which are also shaped by other ecological and physiological constraints.

Overall, our results demonstrate that climate change factors exert strong filtering effects on plant trait coordination, although the nature of these responses varies across species. Previous work has highlighted increased trait coordination along environmental gradients such as rainfall (Li *et al*., 2018) and salinity (Bricca *et al*., 2023). The novelty of our study lies in examining these dynamics within a controlled environment across multiple climate change drivers, offering mechanistic insights into species-specific responses to future climate scenarios.

### Conclusions

Our study highlights the significant impact of individual and interactive effects of climate change factors, including CO_2_, temperature, and drought, on the floral economic traits of *L. corniculatus* and *C. capillaris*. Drought emerged as the dominant factor influencing most plant traits, with species-specific variations in response intensity. The interactive effects of these climate change factors also led to varied responses in floral economic traits. While both species exhibited coordinated trait responses under combined treatments, *C. capillaris* did not show a coordinated response to drought, potentially contributing to its more restricted distribution. To gain a comprehensive understanding of resource allocation strategies (trade-offs among traits) in plants under environmental stresses, larger studies encompassing a broader range of functional traits and larger species set are essential. Investigating the floral economic spectrum and its integration into the whole plant economic spectrum can provide valuable insights into the effects of divergent selective forces on different plant tissues and the potential consequences of shifts in these selective forces as a result of climate change.

## Authors Contributions

Murugash Manavalan: Conceptualization (equal); Investigation (equal), Data curation, Methodology (equal), Formal analysis (lead), Visualization (lead), Writing – original draft (lead), Writing – review and editing (equal); Dinesh Thakur: Conceptualization (equal), Investigation (equal), Methodology (equal), Visualization (supporting), Formal analysis (supporting), Writing – Review & Editing (equal), Supervision (supporting); Michael Bahn: Resources (equal), Funding acquisition for long-term background experiment (lead), Writing – review and editing (supporting); Andreas Schaumberger: Resources (equal), Writing – review and editing (supporting). Zuzana Münzbergová: Conceptualization (equal), Visualization (supporting), Formal analysis (supporting), Resources (equal), Writing – Review & Editing (equal), Supervision (lead).

## AI Assistance

AI assistants, including ChatGPT and Copilot, were utilized to aid in literature gathering, assist with statistical analysis, and refine the tone of the Introduction and Discussion sections. All outputs were meticulously cross-checked by the authors to ensure accuracy and prevent the inclusion of false information in the manuscript.

## Availability of Data and Materials

The data that supports the findings of this study will be made openly available in a Zenodo repository upon acceptance along with the relevant software scripts being attached in GitHub.

## Supplementary Information

Supplementary information has been included in a separate file and detail the following. **Fig. S1.** PCA plots showing trait coordination patterns of *L. corniculatus* under different treatments. **Fig. S2.** PCA plots showing trait coordination patterns of *C. capillaris* under different treatments (Without Seed Traits). **Fig. S3.** PCA plots showing trait coordination patterns of *C. capillaris* under different treatments (With Seed Traits). **Table S1. Number of individuals and plots from which samples of each treatment of *L. corniculatus* and *C. capillaris* were collected. Table S2.** The effect of climate on trait composition tested using RDA on *L. corniculatus* and *C. capillaris*. **Table S3.** Trait degree (number of edges) and betweenness centrality in the network, weighted by correlation strength. **Table S4.** Trait loadings and percentage of trait variation explained by successive principal components (PC) in *L. corniculatus*. **Table S5.** Trait loadings and percentage of trait variation explained by successive principal components (PC) in *C. capillaris*. **Table S6.** Trait loadings of traits and significance of successive constrained axes in RDA in *L. corniculatus*. **Table S7.** Trait loadings of traits and significance of successive constrained axes in RDA in *C. capillaris.* **Table S8.** Dates of induced drought at ClimGrass experimental plots. **Table S9**. Ecological Significance of Studied Plant Traits.

## Acknowledgements

We would like to thank the team of Austrian Research and Education Centre Raumberg-Gumpenstein (HBLFA) for their support during the sampling campaign and for the provision of the experimental site, which was supported by the DaFNE project ClimGrassEco (101067). In addition, we thank N. Rathore and T. Bhatt for assisting with the sampling, as well as D. Parysová for help with the sample processing. The study has been supported by the Czech Science Foundation (project no. 22-00761S) and partly by a long-term research development project no. RVO 67985939 of the Czech Academy of Sciences and institutional support for science and research of the Ministry of Education, Youth, and Sports of the Czech Republic. We thank the POPEKOL discussion group for useful comments on the manuscript.

---

**Table 1.**
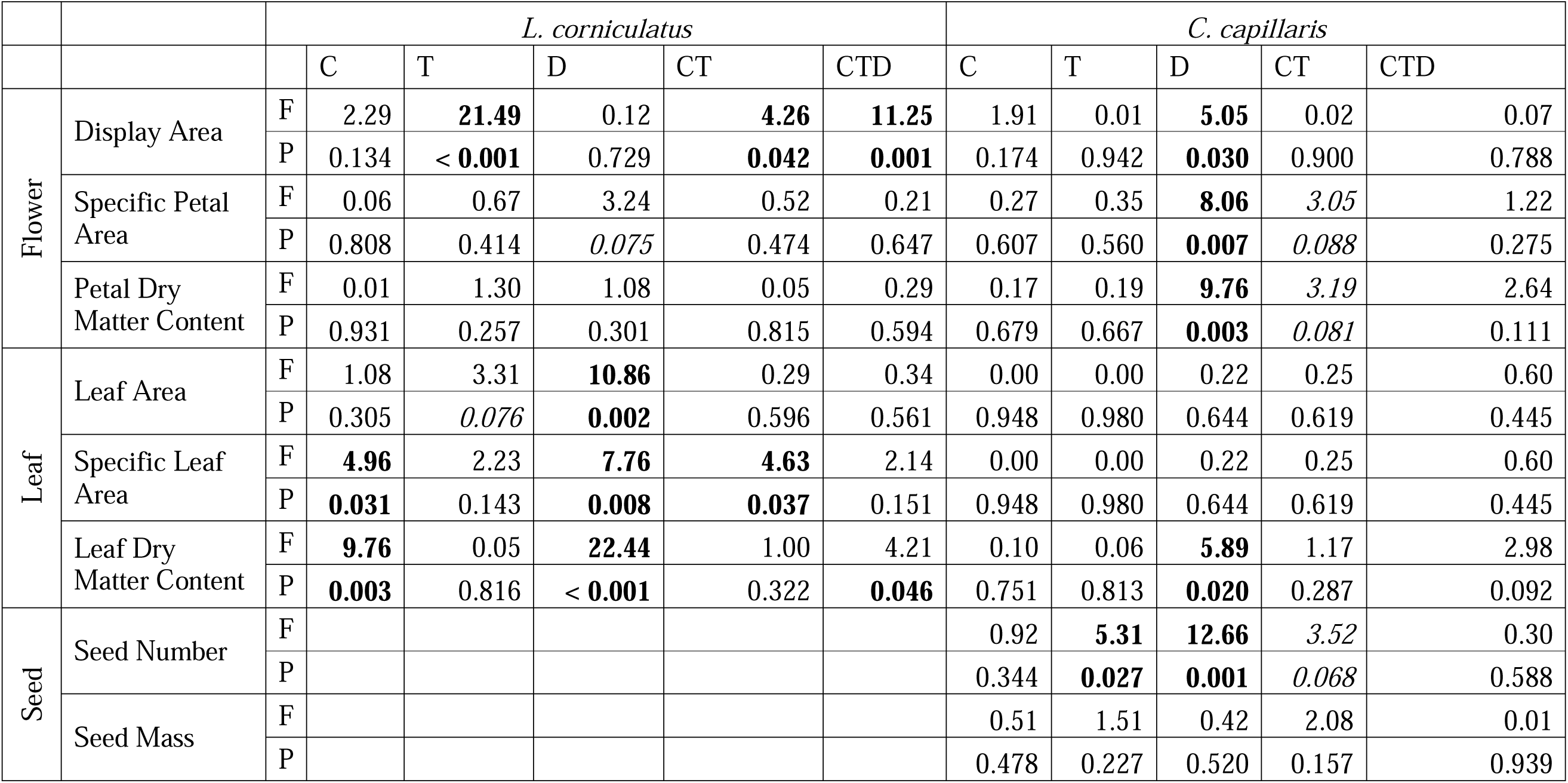
Effect of climate change factors (CO_2_, temperature and drought) on plant traits of *L. corniculatus* and *C. capillaris.* Significant effects values < 0.05) are marked in bold while marginally significant effects (p-values between 0.05 and 0.1) are marked in italics

|  |  | *L. corniculatus* |  |  |  |  |  | *C. capillaris* |  |  |  |  |
| --- | --- | --- | --- | --- | --- | --- | --- | --- | --- | --- | --- | --- |
|  |  |  | C | T | D | CT | CTD | C | T | D | CT | CTD |
| Flower | Display Area | F | 2.29 | **21.49** | 0.12 | **4.26** | **11.25** | 1.91 | 0.01 | **5.05** | 0.02 | 0.07 |
|  |  | P | 0.134 | < **0.001** | 0.729 | **0.042** | **0.001** | 0.174 | 0.942 | **0.030** | 0.900 | 0.788 |
|  | Specific Petal Area | F | 0.06 | 0.67 | 3.24 | 0.52 | 0.21 | 0.27 | 0.35 | **8.06** | *3.05* | 1.22 |
|  |  | P | 0.808 | 0.414 | *0.075* | 0.474 | 0.647 | 0.607 | 0.560 | **0.007** | *0.088* | 0.275 |
|  | Petal Dry Matter Content | F | 0.01 | 1.30 | 1.08 | 0.05 | 0.29 | 0.17 | 0.19 | **9.76** | *3.19* | 2.64 |
|  |  | P | 0.931 | 0.257 | 0.301 | 0.815 | 0.594 | 0.679 | 0.667 | **0.003** | *0.081* | 0.111 |
| Leaf | Leaf Area | F | 1.08 | 3.31 | **10.86** | 0.29 | 0.34 | 0.00 | 0.00 | 0.22 | 0.25 | 0.60 |
|  |  | P | 0.305 | *0.076* | **0.002** | 0.596 | 0.561 | 0.948 | 0.980 | 0.644 | 0.619 | 0.445 |
|  | Specific Leaf Area | F | **4.96** | 2.23 | **7.76** | **4.63** | 2.14 | 0.00 | 0.00 | 0.22 | 0.25 | 0.60 |
|  |  | P | **0.031** | 0.143 | **0.008** | **0.037** | 0.151 | 0.948 | 0.980 | 0.644 | 0.619 | 0.445 |
|  | Leaf Dry Matter Content | F | **9.76** | 0.05 | **22.44** | 1.00 | 4.21 | 0.10 | 0.06 | **5.89** | 1.17 | 2.98 |
|  |  | P | **0.003** | 0.816 | < **0.001** | 0.322 | **0.046** | 0.751 | 0.813 | **0.020** | 0.287 | 0.092 |
| Seed | Seed Number | F |  |  |  |  |  | 0.92 | **5.31** | **12.66** | 3.52 | 0.30 |
|  |  | P |  |  |  |  |  | 0.344 | **0.027** | **0.001** | *0.068* | 0.588 |
|  | Seed Mass | F |  |  |  |  |  | 0.51 | 1.51 | 0.42 | 2.08 | 0.01 |
|  |  | P |  |  |  |  |  | 0.478 | 0.227 | 0.520 | 0.157 | 0.939 |

