## Supplementary material for "Elevated CO2, Warming and Drought Differentially Impact Reproductive and Vegetative Economic Traits in Two Grassland Species": Supplmentary Information

**Supplementary Information**

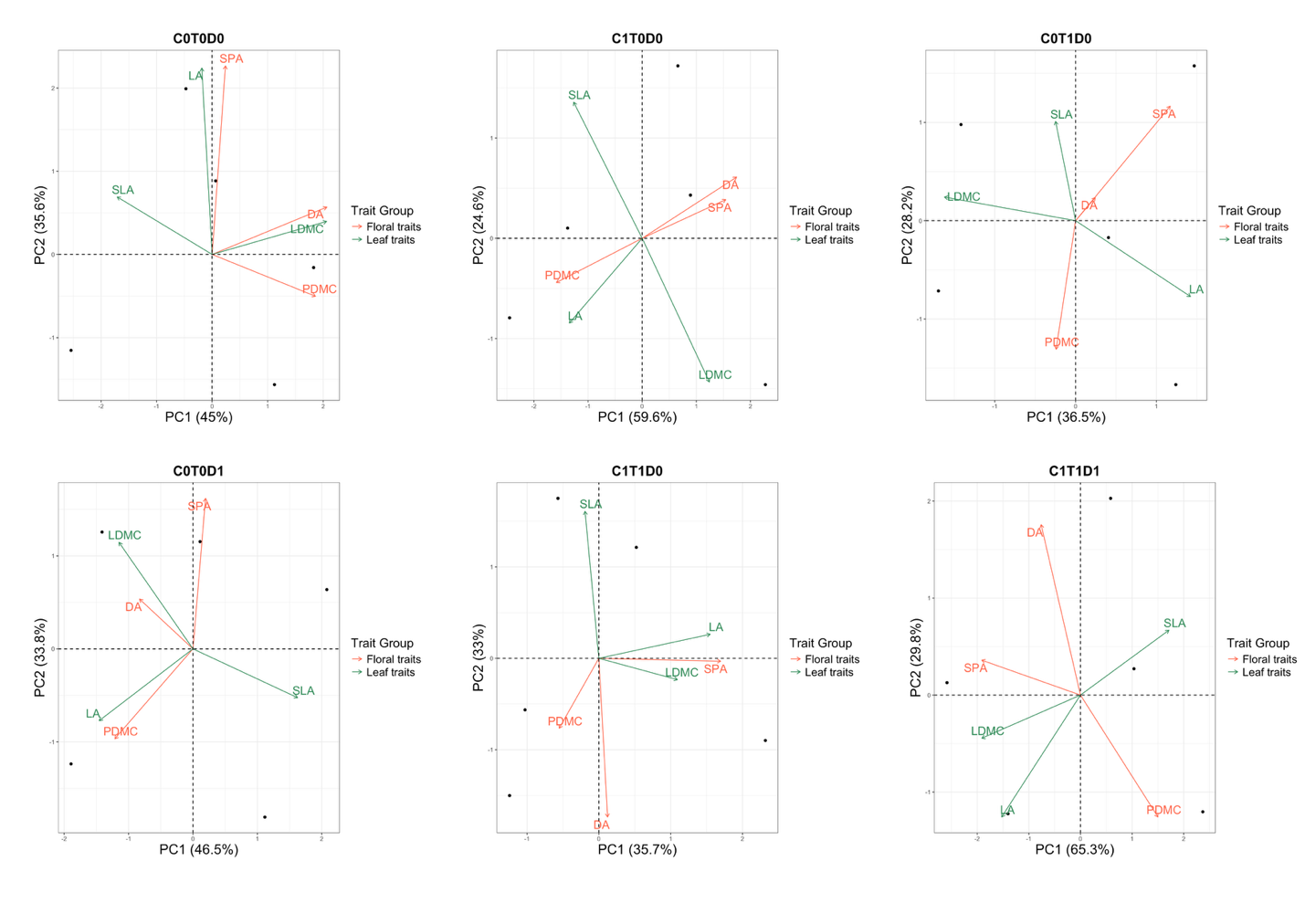

**Fig. S1.** PCA plots showing trait coordination patterns of *L. corniculatus* under different treatments. C0T0D0 - Ambient Control, C1T0D0 – Elevated CO_2_ (+300ppm), C0T1D0 – Warming (+3°C), C0T0D1 – Drought, C1T1D0 – Elevated CO_2_ combined with warming, C1T1D1 – Elevated CO_2_, warming and drought.

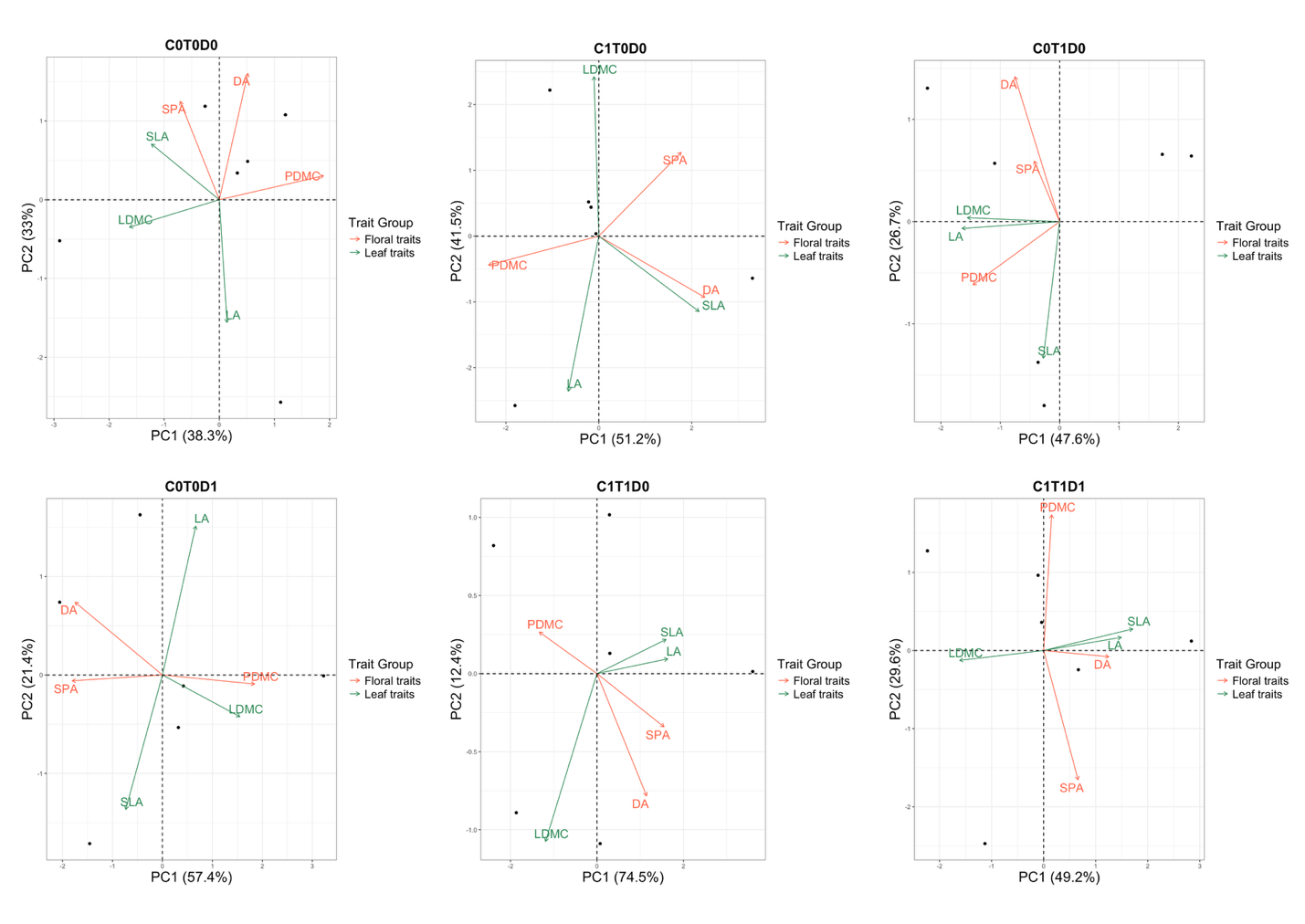

**Fig. S2.** PCA plots showing trait coordination patterns of *C. capillaris* under different treatments (Without Seed Traits). C0T0D0 - Ambient Control, C1T0D0 – Elevated CO_2_ (+300ppm), C0T1D0 – Warming (+3°C), C0T0D1 – Drought, C1T1D0 – Elevated CO_2_ combined with warming, C1T1D1 – Elevated CO_2_, warming and drought.

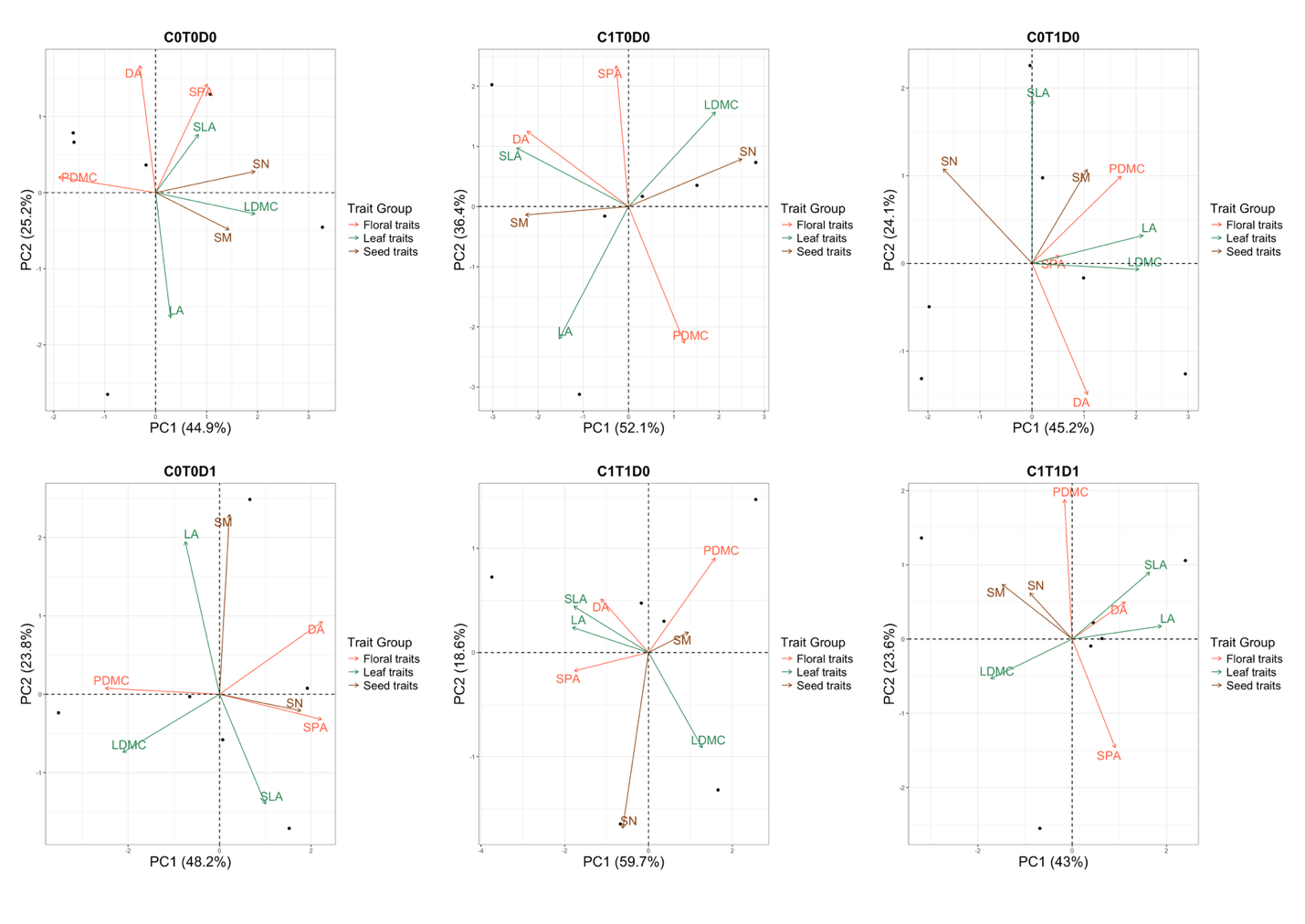

**Fig. S3.** PCA plots showing trait coordination patterns of *C. capillaris* under different treatments (With Seed Traits). C0T0D0 - Ambient Control, C1T0D0 – Elevated CO_2_ (+300ppm), C0T1D0 – Warming (+3°C), C0T0D1 – Drought, C1T1D0 – Elevated CO_2_ combined with warming, C1T1D1 – Elevated CO_2_, warming and drought.

|  | *Lotus corniculatus* L. | | *Crepis capillaris* L. | |
| --- | --- | --- | --- | --- |
| Treatment | No. of Individuals | No. of Plots | No. of Individuals | No. of Plots |
| C0T0D0 | 11 | 8 | 8 | 4 |
| C2T0D0 | 6 | 4 | 6 | 3 |
| C0T2D0 | 6 | 4 | 10 | 4 |
| C2T2D0 | 5 | 4 | 7 | 3 |
| C0T0D1 | 5 | 4 | 8 | 4 |
| C2T2D1 | 6 | 4 | 6 | 3 |
| **Total** | **39** | **28** | **45** | **21** |

**Table S1.** Number of individuals and plots from which samples of each treatment of *L. corniculatus* and *C. capillaris* were collected.

|  | *L. corniculatus* | | *C. capillaris* | |
| --- | --- | --- | --- | --- |
|  | Variance (%) | P-value | Variance (%) | P-value |
| CO2 | 4.1 | 0.105 | 2.1 | 0.420 |
| Temperature | 6.3 | **0.022** | 2.9 | 0.196 |
| Drought | 10.1 | **0.002** | 7.7 | **0.001** |
| C : T | 1.7 | 0.543 | 4.5 | **0.037** |
| (C + T) : D | 4.8 | *0.070* | 2.8 | 0.206 |
| Residual Variance | 72.9 |  | 80.0 |  |

**Table S2.** The effect of climate on trait composition tested using RDA on *L. corniculatus* and *C. capillaris*

|  | *L. corniculatus* | | *C. capillaris* | |
| --- | --- | --- | --- | --- |
| Trait | Weighted  degree | Weighted betweenness | Weighted  degree | Weighted betweenness |
| DA | 0.39 | 0.00 | 0.47 | 0.00 |
| SPA | 0.55 | 0.00 | 1.08 | 0.14 |
| PDMC | 0.94 | 0.10 | 1.06 | 0.19 |
| LA | 0.00 | 0.00 | 0.00 | 0.00 |
| SLA | 0.76 | 0.00 | 0.00 | 0.00 |
| LDMC | 0.76 | 0.00 | 0.38 | 0.00 |
| SN |  |  | 0.83 | 0.14 |
| SM |  |  | 0.00 | 0.00 |

**Table S3.** Trait degree (number of edges) and betweenness centrality in the network, weighted by correlation strength.

| *L. corniculatus* | | | | | | |
| --- | --- | --- | --- | --- | --- | --- |
|  | PC1 | PC2 | PC3 | PC4 | PC5 | PC6 |
| DA | -0.40 | 0.31 | **-0.24** | **0.81** | -0.16 | -0.08 |
| SPA | -0.10 | **0.62** | 0.33 | **-0.31** | **-0.63** | 0.00 |
| PDMC | 0.31 | **-0.57** | 0.09 | 0.26 | **-0.70** | 0.13 |
| LA | 0.37 | 0.16 | **0.77** | **0.41** | 0.29 | 0.03 |
| SLA | **-0.59** | -0.21 | 0.30 | -0.04 | 0.07 | **0.72** |
| LDMC | **0.50** | 0.36 | -0.39 | 0.07 | 0.03 | 0.68 |
| Cumulative (%) | 35.24 | 64.06 | 79.00 | 90.22 | 96.39 | 100 |

**Table S4.** Trait loadings and percentage of trait variation explained by successive principal components (PC) in *L. corniculatus*.

| *C. capillaris* | | | | | | | | |
| --- | --- | --- | --- | --- | --- | --- | --- | --- |
|  | PC1 | PC2 | PC3 | PC4 | PC5 | PC6 | PC7 | PC8 |
| DA | **0.44** | **-0.74** | 0.06 | **-0.50** | **-0.54** | **0.73** | **0.55** | 0.35 |
| SPA | **0.54** | **-0.59** | -0.22 | -0.08 | 0.09 | -0.43 | -0.12 | -0.76 |
| PDMC | **-0.62** | 0.10 | 0.09 | -0.19 | **-0.40** | 0.06 | **0.41** | -0.67 |
| LA | 0.08 | 0.07 | **0.62** | **-0.55** | 0.09 | **-0.42** | 0.04 | 0.11 |
| SLA | 0.25 | 0.33 | -0.34 | -0.01 | **-0.46** | **-0.41** | 0.25 | 0.17 |
| LDMC | -0.32 | -0.35 | -0.18 | 0.07 | 0.31 | -0.31 | **0.43** | 0.25 |
| SN | 0.29 | 0.24 | 0.15 | 0.19 | 0.28 | 0.14 | **0.44** | -0.16 |
| SM | 0.00 | -0.12 | 0.23 | 0.29 | -0.17 | -0.08 | 0.00 | 0.02 |
| Cumulative (%) | 27.95 | 46.56 | 61.81 | 73.03 | 82.44 | 91.19 | 97.70 | 100.00 |

**Table S5.** Trait loadings and percentage of trait variation explained by successive principal components (PC) in *C. capillaris.*

| *L. corniculatus* | | | | | | |
| --- | --- | --- | --- | --- | --- | --- |
|  | RDA 1 | RDA 2 | RDA 3 | RDA 4 | RDA 5 | PC 1 |
| DA | -0.61 | -0.56 | -0.41 | 0.03 | -0.14 | 0.76 |
| SPA | -0.16 | 0.24 | 0.28 | 0.23 | -0.14 | 1.29 |
| PDMC | 0.21 | 0.24 | -0.20 | -0.25 | 0.00 | -1.29 |
| LA | -0.44 | 0.67 | -0.43 | 0.10 | 0.00 | 0.08 |
| SLA | -0.95 | -0.12 | 0.04 | 0.07 | 0.23 | -0.16 |
| LDMC | 1.05 | -0.16 | -0.29 | 0.21 | 0.10 | 0.47 |
| Pr(>F) | **0.0006** | 0.5574 | 0.9501 | 0.9767 | 0.9767 | NA |
| Cumulative (%) | 17.44 | 23.49 | 27.19 | 28.33 | 29.00 | 56.64 |

**Table S6.** Trait loadings of traits and significance of successive constrained axes in RDA in *L. corniculatus*. Significant Climatic Factors: Drought (p = 0.0008) and Temperature (p = 0.0223).

| *C. capillaris* | | | | | | |
| --- | --- | --- | --- | --- | --- | --- |
|  | RDA 1 | RDA 2 | RDA 3 | RDA 4 | RDA 5 | PC 1 |
| DA | -0.37 | -0.16 | 0.22 | -0.12 | -0.01 | 1.04 |
| SPA | -0.53 | -0.06 | -0.11 | 0.07 | -0.06 | 1.26 |
| PDMC | 0.77 | -0.21 | 0.17 | 0.08 | -0.02 | -0.76 |
| LA | -0.06 | -0.66 | 0.10 | 0.11 | -0.05 | -0.03 |
| SLA | -0.54 | 0.19 | 0.16 | 0.34 | 0.04 | -0.26 |
| LDMC | 0.65 | -0.29 | -0.22 | 0.09 | 0.08 | 0.28 |
| SN | -0.83 | -0.38 | -0.07 | -0.10 | 0.07 | -0.20 |
| SM | 0.17 | 0.02 | 0.39 | -0.08 | 0.05 | 0.49 |
| Pr(>F) | **0.0005** | 0.5049 | 0.9673 | 0.9906 | 0.9998 | NA |
| Cumulative (%) | 13.04 | 17.15 | 18.94 | 19.90 | 20.02 | 39.56 |

**Table S7.** Trait loadings of traits and significance of successive constrained axes in RDA in *C. capillaris.* Significant Global Change Factors: Drought (p = 0.0012) and C×T (p = 0.0395).

| **Year** | **Growth Period** | **Drought Dates** |
| --- | --- | --- |
| 2017 | 2nd growth after 1st cut | 31.05.2017 – 26.07.2017 |
| 2019 | 1st growth from start of growing season | 18.04.2019 – 05.06.2019 |
| 2020 | 2nd growth after 1st cut | 17.06.2020 – 29.07.2020 |
| 2021 | 2nd growth after 1st cut | 27.05.2021 – 28.07.2021 |
| 2022 | 2nd growth after 1st cut | 25.05.2022 – 27.07.2022 |
| 2023 | 2nd growth after 1st cut | 24.05.2023 – 26.07.2023 |
| 2024 | 2nd growth after 1st cut | 29.05.2024 – 19.08.2024 |

**Table S8.** Dates of induced drought at ClimGrass experimental plots.

| Trait | Ecological Significance | Reference |
| --- | --- | --- |
| Display Area (DA) | It serves as a visual signal that enhances pollinator attraction. Larger displays increase the probability of visitation, which can lead to higher pollination success, seed set, and reproductive output. Beyond attraction, display area may also influence pollinator behavior, such as visit duration and flower constancy. However, larger displays can come with costs, including increased resource investment and exposure to herbivores or abiotic stress (e.g., wind, desiccation). | (Conner and Rush, 1996; Delgado *et al.*, 2023) |
| Specific Petal Area (SPA) | Construction cost per unit area of petals. Low SPA generally means thicker petals which reflects a shift towards less economical flowers, implying higher construction costs per unit area. It has been found to be negatively correlated with corolla lifespan and with water allocation. | (Roddy, Brodersen and Dawson, 2016; Zhang *et al.*, 2017) |
| Petal Dry Matter Content (PDMC) | Indicates tissue density and structural investment per unit fresh mass. Lower PDMC can be associated with less denser, more economical tissues that are cheaper to produce and are less resilient to stress. It positively relates to flower longevity. | (Roddy *et al.*, 2023) |
| Leaf Area (LA) | Plays a critical role in light interception, influencing photosynthetic capacity and energy acquisition. Larger leaves can enhance competitive ability for light, especially in dense plant communities or shaded environments. However, they may also increase water loss and be more vulnerable to heat or drought stress. | (Díaz *et al.*, 2016; Li and Prentice, 2024) |
| Specific Leaf Area (SLA) | A key trait in the leaf economics spectrum reflecting leaf construction cost per unit area. High SLA is typically associated with fast-growing species that invest in thin, resource-efficient leaves, enabling rapid photosynthesis and growth, especially in resource-rich or disturbed environments. | (Wright *et al.*, 2004; Díaz *et al.*, 2016) |
| Leaf Dry Matter Content (LDMC) | Another key trait in leaf economics, reflecting how much dry mass a leaf contains relative to its fresh weight. High LDMC is associated with tougher, longer-lived leaves that are more resistant to environmental stress. | (Pakeman, Eastwood and Scobie, 2011; Blumenthal *et al.*, 2020) |
| Seed Mass (SM) | Seed Mass plays a crucial ecological role by influencing plant dispersal strategies, establishment success, and adaptation to environmental conditions. Heavier diaspores are often associated with coarser root systems, which may enhance seedling survival in challenging habitats but can limit dispersal distance, making these species more reliant on local establishment rather than long-range colonization. | (Díaz *et al.*, 2016; Bergmann *et al.*, 2017; Li and Prentice, 2024) |
| Seed Number (SN) | Key fitness estimate. High seed number is associated with colonization ability and reproductive output, especially in disturbed or variable environments. Often inversely related to seed mass. | (Leishman, 2001; Henrik Bruun and Poschlod, 2006) |

**Table S9.** Ecological Significance of Studied Plant Traits.
